# Structural Connectivity Between the Zinc Linchpin Motif, the [4Fe-4S] cluster, and the active site orchestrates DNA repair in MUTYH

**DOI:** 10.64898/2026.05.27.728240

**Authors:** Mohammad Hashemian, Melody Malek, Tian Xia, April Lu, Alam X. León-Cardona, Andrew Fisher, Carlos H. Trasviña-Arenas, Sheila S. David

**Affiliations:** Department of Chemistry, University of California, Davis, California, 95616, United States; Graduate Program in Chemistry and Chemical Biology, University of California, Davis, CA 95616, USA; Department of Molecular and Cellular Biology, University of California, Davis, California 95616, United States; Centro de Investigación sobre el Envejecimiento, Centro de Investigación y de Estudios Avanzados (CINVESTAV), Unidad Sede Sur, Tlalpan, Ciudad de México 14330, Mexico

**Author notes:** Corresponding authors: Carlos H. Trasvina-Arenas, Sheila S. David.

**Keywords:** MutY, Base Excision Repair, DNA glycosylase, [4Fe-4S] cluster, Zinc Linchpin Motif, allostery

## Abstract

The DNA glycosylase MUTYH protects genomic integrity by excising adenine mispaired with 8-oxoguanine (OG), which initiates base excision repair (BER). The [4Fe-4S] cluster DNA binding domain and Zn linchpin motif of MUTYH are hotspots for inherited cancer associated variants (CAVs) highlighting their critical functions in DNA repair. Here, we present three full-length human MUTYH crystal structures bound to DNA across three catalytic states, representing early and late transition states and product complexes. These structures reveal a previously unrecognized interplay between the Zn site and the Fe–S cluster mediated by His85 and a conserved Arg247/Arg307 bridge spanning ∼15 Å. Disruption of this network impairs metal loading, reduces the active enzyme fraction, weakens lesion DNA binding, and diminishes OG:A repair in cells. Moreover, the Zn site adopts distinct coordination modes across the reaction coordinate, with a water molecule replacing a cysteine ligand in transition state analog complexes and cysteine coordination restored in a product-mimicked state. Functional analyses show that this ligand switching is dispensable for core glycosylase chemistry *in vitro* but disproportionately affects repair in cells, suggesting an additional role of the Zn linchpin in cellular OG:A repair beyond intrinsic glycosylase activity, likely involving interactions with BER partners. Together, these findings illustrate how the Zn linchpin enhances Fe–S cluster domain DNA substrate engagement to enable effective DNA repair and provides a rationale for how these functions are compromised by MUTYH CAVs.

## INTRODUCTION

Reactive oxygen and nitrogen species (RONS), generated endogenously during cellular metabolism or introduced from environmental sources, inflict oxidative damage on DNA, producing a wide array of modified nucleobases that threaten genomic integrity. The base excision repair (BER) pathway has evolved to recognize and remove oxidized bases, thereby restoring correct genetic information.^1–5^ 8-oxoguanine (OG) is the most prevalent oxidized nucleobase,^6^ and its ability to mispair with adenine to form the promutagenic OG:A base pair drives the accumulation of genomic G:C→T:A transversion mutations. The human MutY homolog (MUTYH) is unique among BER glycosylases in its exclusive role in removing adenine opposite OG, thereby enabling the OG glycosylase (OGG1) and downstream BER enzymes to restore the canonical G:C base pair.^7^ Inherited MUTYH variants with compromised activity lead to accumulation of G→T transversion mutations and is linked to an increased predisposition to colorectal cancer in a hereditary syndrome known as MUTYH-associated polyposis (MAP). ^8–14^

MutY glycosylases are multi-domain enzymes that belong to the N-terminal Helix-hairpin-Helix (HhH) family with a catalytic domain that harbors the active site residues and a conserved [4Fe-4S]^2^ cluster^10,15–20^ and a C-terminal OG-recognition domain that confers specificity for 8-oxoguanine mis-paired with the target adenine^21–26^. A flexible interdomain connector (IDC) connects the N- and C-terminal domains and in the human protein is proposed to serve as a protein-protein interaction platform with downstream BER, checkpoint signaling, and chromatin remodeling factors. ^27–29,9,30,31^. In humans and other Metazoa, a structurally expanded IDC introduces a second metal cofactor through the emergence of a Zn coordinating ‘linchpin’ motif.^30,32,33^ Conspicuously, many cancer-associated MUTYH “variants of uncertain significance” (VUS) are localized around two of the protein’s metallocofactors.^9,18^

The [4Fe-4S] cluster and its associated iron-sulfur cluster loop (FCL) in MUTYH are required for substrate DNA binding and proper engagement of the lesion base pair to support adenine excision.^34–40^ In recent work, we showed that the [4Fe-4S] cluster serves as an allosteric site, facilitating catalysis through a conserved network of polar contacts, maintained even among ‘cluster-less’ MutY homologs, where alternative secondary structural elements arise to preserve this allosteric bridge. ^10,14,41,42^ Consistent with its roles in lesion engagement and allostery, MUTYH cancer-associated variants (CAVs) surrounding the [4Fe-4S] cluster compromise glycosylase activity. Remarkably, even variants that *do not* result in cluster loss but *do* disrupt inter-side chain hydrogen bonding within the allosteric bridge to the active site still exhibit complete loss of glycosylase activity.^10,20^ Overall, the [4Fe-4S] cluster in MUTYH serves multiple, well-defined biochemical roles that are more clearly established than those of the Zn site.

Previous work from our laboratory indicated that MUTYH, alongside other Metazoan MutY homologs and in contrast to bacterial and archaeal MutY enzymes, has acquired a second metal-binding site in addition to the [4Fe-4S] cluster conserved across all three domains of life. In this site, Zn coordination was hypothesized to form a ‘linchpin’, reinforcing engagement of the two functional domains of MUTYH to facilitate accurate DNA lesion repair.^32,33^ Subsequent structural analysis of mouse Mutyh proposed a 3-Cys/1-His coordination environment (corresponding to residues His85, Cys332, Cys339, and Cys342 in the *human* homolog, MUTYH), yet the available mouse structure shares only ≈60% sequence identity with human MUTYH’s IDC (and 70% global sequence identity) and lacks structural resolution of much of the IDC, limiting its ability to fully represent the human Zn 3D motif and surrounding cancer-associated variant residues.^30^ Furthermore, a more recent high-resolution human MUTYH-DNA complex is devoid of the Zn cofactor, leaving the architecture of the human Zn-binding motif unresolved.^10^ Nonetheless, biochemical and mutational studies consistently indicate that disruption of predicted Zn-ligating residues diminishes OG:A binding and glycosylase activity, underscoring the functional importance of the Zn cofactor.^32,33^ Together, these findings show that, although far less defined than the [4Fe-4S] cluster, the Zn linchpin is emerging as an auxiliary feature assisting in proper MUTYH engagement of damaged DNA during repair.

In the current work, we present a set of full-length human MUTYH structures bound to DNA duplexes, forming three distinct complexes that retain both metal cofactors, thereby resolving the architecture of the Zn-binding linchpin and its relationship to the [4Fe-4S] cluster. These structures reveal an alternative and active site state-dependent Zn coordination environment. In the early transition state analog complex (TSAC 1), Zn coordination by Cys332 is observed in only one chain, while a water molecule occupies this position in the other chain; in the late transition state analog complex (TSAC 2), this position is occupied by water in both chains. Coordination of Cys332, in both chains, is only observed in the product-analog complex (PAC). Our findings suggest that Zn loading is associated with a higher fraction of active enzyme, tighter OG:A binding, enhanced [4Fe-4S] cluster loading, improved OG:A repair in cells, and a stronger association with the downstream AP endonuclease (APE1), as assessed by glycosylase assays, ICP-MS, cellular repair assays, and AlphaFold-Multimer predictions. Finally, comparative structural and evolutionary analyses identified two arginine residues (Arg247 and Arg307) that co-emerged with the Zn site in higher eukaryotes and form a conserved bridge linking the Zn center to both the [4Fe-4S] cluster and active site of MUTYH. Disrupting this bridge attenuates glycosylase activity and OG:A repair in cells, highlighting an unanticipated layer of allosteric communication that is essential for optimal MUTYH function. Moreover, these findings may inform functional impacts of MUTYH CAVs that exist within this broad network connecting both metal sites and the catalytic pocket, as well as those which reside on the opposite, solvent exposed face of the Zn center implicated in handoff of the AP-site containing product to downstream BER partners.

## METHODS

### *MUTYH* gene cloning and mutagenesis

MUTYH proteins were purified following methods previously described.^10^ For crystallographic studies, the construct was further truncated by deletion of the first 34 and final 28 codons, analogous to truncations used in the murine MUTYH structure.^30^ The primary amino acid sequence of MUTYH used for crystallography is depicted in **Figure S1.** Site-directed mutagenesis of pET28-MBP-MUTYH was performed by PCR-based overlap extension and in vivo assembly to generate TOP10 clones bearing the mutated plasmid.^43,44^

### MUTYH overexpression and purification

MUTYH overexpression and purification were adapted from previously reported protocols with modifications to preserve native metal cofactors.^10,26^ A BL21(DE3) strain harboring pRKISC and pKJE7 plasmids – coexpressing the [4Fe–4S] cluster assembly machinery^33^ and the DnaK/DnaJ/GrpE chaperone system^45^ – was transformed with pET28-MBP-MUTYH and selected on LB agar containing kanamycin, tetracycline, and chloramphenicol. Cultures were grown in Terrific Broth to an OD_600_ ≥1.5 at 37 °C with shaking at 180 rpm, cooled for 1 h at 4 °C, and induced with IPTG (1 mM) in the presence of ferrous sulfate and ferric citrate (at a final concentration of 50mg/L culture) to support [4Fe-4S] cluster assembly; ZnSO_4_ (at a final concentration of 60mg/L culture) supplementation was delayed until ≈20 h post-induction to avoid inhibition of Fe–S cluster biosynthesis and maturation pathways^46,47^ **(Figure S2)**, and cells were harvested 2 hours after supplementation with ZnSO_4_ for a total induction time of 22 hours at 15 °C with shaking at 180 rpm. All subsequent purification steps were conducted either using EDTA-free buffers to prevent Zn chelation for generation of MUTYH(Zn+) proteins *or* with 2mM EDTA for generation of MUTYH(Zn-) (Zn deficient) protein. Cell pellets were resuspended in lysis buffer (30 mM Tris–HCl, pH 7.5, 1 M NaCl, 30 mM β-mercaptoethanol, 10% glycerol) supplemented with PMSF (1 mM), lysed by sonication on ice, and clarified by centrifugation. The clarified supernatant was filtered and incubated with Ni^2+^ NTA resin and washed extensively with lysis buffer prior to elution with buffer containing 30 mM Tris HCl (pH 7.5), 200 mM NaCl, 30 mM β-mercaptoethanol, 10% glycerol, and 500 mM imidazole. Buffer exchange and further purification were performed by heparin-affinity chromatography using buffer A (30 mM Tris–HCl, pH 7.5, 10% glycerol, 1 mM DTT) and buffer B (30 mM Tris–HCl, pH 7.5, 1 M NaCl, 10% glycerol, 1 mM DTT), followed by concentration and analysis by SDS–PAGE. Protein concentration was determined by UV absorbance at 280 nm with a predicted extinction coefficient of 84,630 M^-1^. MBP and His tags were removed by TEV protease digestion following nickel-affinity chromatography, and the protein was further purified by a second heparin-affinity step and, for crystallography, an additional size-exclusion chromatography step using 20% buffer B on a Superdex 200 Increase 10/300 GL column. An SDS PAGE gel of all purified proteins represented in this work is shown in **Figure S4B.**

### Preparation of oligonucleotide substrates

The 1N and 1NBn transition state analog phosphoramidites were synthesized in house with modifications of previously reported procedures.^48^ These modified phosphoramidites, along with OG-deoxyribonucleotides, were incorporated into oligonucleotides by solid-phase DNA synthesis. Synthesized DNA strands were cleaved from solid support and deprotected using ammonium hydroxide supplemented with 0.25 M β-mercaptoethanol for 17 h at 55 °C. All oligonucleotides were purified by HPLC or gel purification using 13% (for 11 nt DNAs) or 19% (for 30 nt DNAs) denaturing PAGE gels, desalted using Sep-Pak C18 cartridges (Waters), and verified by MALDI mass spectrometry at the UC Davis Campus Mass Spectrometry Facility. The tetrahydrofuran (THF), and 2’fluoro-2’deoxyadenosine (fA)-containing oligonucleotides were purchased from Integrated DNA Technologies (IDT) and, together with the in-house–synthesized 1N- and 1NBn-containing oligonucleotides, were annealed to complementary OG-containing oligonucleotides for biochemical or crystallographic studies. For kinetic and EMSA analyses, the 30-nt adenine strand was 5’-end radiolabeled with [γ-^32^P]ATP using T4 polynucleotide kinase (PNK) before annealing to the complementary OG-containing strand such that 5% of the duplex DNA was labeled for kinetics and 100% labeled for EMSA.

**Table 1.**
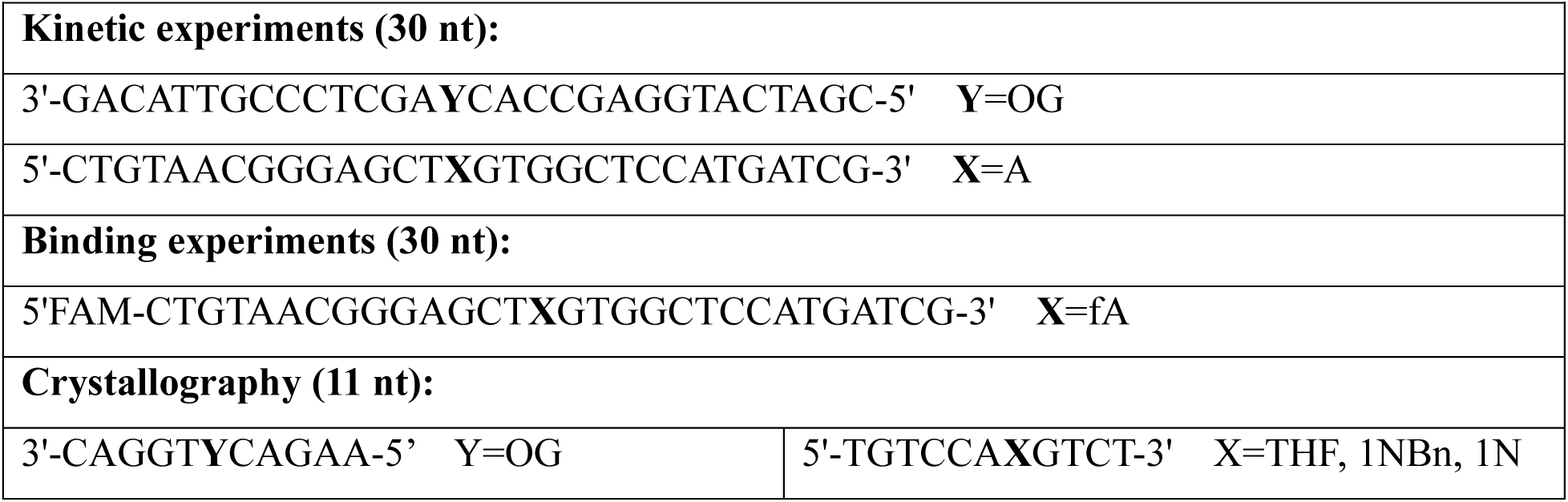
Oligonucleotides used in this study.

### MUTYH crystallography

Crystallization conditions for human MUTYH were based on previously reported conditions for related protein-DNA complexes.^10,30^ The 1N/1NBn/THF and OG-containing oligonucleotides used for crystallography were annealed at a 1:1 molar ratio (263 µM) in 30 mM Tris–HCl (pH 7.5) by heating to 90 °C for 5 min followed by slow cooling to 4 °C. The resulting duplexes (263 µM) were mixed with MUTYH protein (236 µM, 12.6 mg/ml) in buffer containing 30 mM Tris–HCl (pH 7.5), 100 mM NaCl, and 0.5 mM DTT to form the MUTYH–DNA complex (final concentration 131 µM). After incubating at room temperature for 30 min, the complex was combined with crystallization solution in a 1:1 ratio and set up as 3 µL hanging drops on coverslips (Hampton Research). Crystals were grown at room temperature by hanging-drop vapor diffusion, with optimal crystals obtained in 0.1 M Bis–Tris (pH 5.5-6.0), 0.2 M ammonium sulfate, and 20% (w/v) PEG 3350. Golden rod- or needle-like crystals appeared within 12–24 h. Crystals were harvested directly from the drop and flash-cooled in liquid nitrogen without additional cryo protectant.

Prior to diffraction data collection, a Zn K-edge X-ray absorption spectrum was collected on the first batch of MUTYH(Zn+) crystals grown in complex with THF:OG-containing duplex DNA to confirm that Zn incorporation was maintained throughout the purification and crystallization workflow **(Figure S3).** X-ray diffraction data were collected using 0.2 ° oscillation at beamline 12-1 at the Stanford Synchrotron Radiation Lightsource (SSRL). Diffraction data were processed using XDS^49^ and scaled with AIMLESS^50^ Initial phases were obtained by molecular replacement using Phaser as implemented in PHENIX^51^ with a previously determined MUTYH-DNA structure^10^ (PDB:8FAY) used as the model. Model refinement was carried out in PHENIX using iterative cycles of simulated annealing, restrained refinement, and manual model building using COOT^52^, including appropriate restraints for the 1N and 1NBn transition state analogs, OG, THF, Zn, and the [4Fe-4S]^2^ cluster. Data collection and refinement statistics are summarized in Supplementary **Table S1**. Structural figures for visualization were generated using PyMOL. Coordinates and structure factors have been deposited in the Protein Data Bank under accession codes:11ji, 11jh and 11ip for MUTYH in complex with 1N:OG, 1NBn:OG and THF:OG-containing DNA duplex, respectively.

### MUTYH–APE1 interaction predictions

Predictive modeling of the MUTYH-APE1-DNA ternary complex was performed using AlphaFold3 Multimer, as implemented in ColabFold, a streamlined and user-accessible AlphaFold workflow.^53,54^. Given the intrinsic nature of MUTYH and APE1 to bind DNA, the length of the DNA sequence was 90 base pairs (bps) to enable sufficient DNA-protein interaction and minimize biasing in MUTYH-APE1 binding. Moreover, the DNA contained a centrally located THF:OG lesion. No additional restraints or interface constraints were applied. The derived five models were analyzed and filtered based on the conformation of active site residues, DNA engagement, and Zn coordination, and the selected model resulted with the highest structural alignment of these regions with the MUTYH-THF:OG crystal structure (PDB ID 11ip; **Figure S10A-C**). The selected model was subjected to angle and bond relaxation following previously described Rossetta protocols.^55,56^

### Glycosylase assay and binding experiments

The glycosylase (*k_2_*) and turnover (*k_3_*) rate constants of **MUTYH** with OG:A-containing 30-bp DNA substrates were measured under single-turnover (STO) and multiple-turnover (MTO) conditions, respectively, as previously described,^17^ following the minimal kinetic scheme:

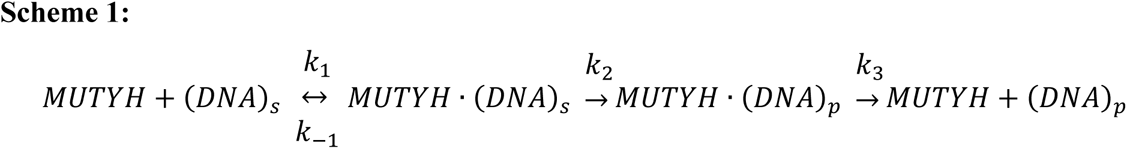

(DNA)*_s_*denotes the OG:A-containing DNA substrate and (DNA)_p_denotes the abasic-site product formed following MUTYH-catalyzed adenine excision. Active enzyme concentrations were determined by active-site titrations (AST) by quantifying the amplitude of the burst phase (A_0_) in biphasic product formation curves collected under MTO conditions using 20 nM total enzyme and 20 nM OG:A substrate at 37 °C. Turnover rate constants (*k_3_*) were determined from the linear steady-state phase of the biphasic curves using the same conditions. Product formation data were fit using the following equations to extract the active fraction and kinetic parameters, where *k_ss_*corresponds to the slope of the linear phase:^17^

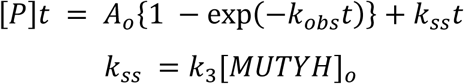

Glycosylase rate constants (k_2_) were measured under STO conditions with 100 nM active enzyme and 20 nM OG:A substrate at 20 °C. Product formation curves were fit to a single exponential function to determine the observed rate constant (k_obs_), which under STO conditions approximates k_2_ (37):

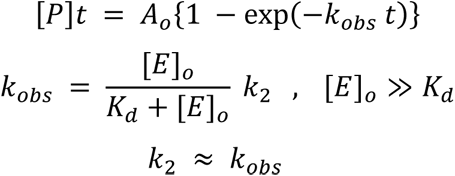

For all glycosylase assays, the reaction composition (excluding enzyme) was as follows: 0.1 mg/mL BSA, 30mM NaCl, 20 mM Tris-HCl pH 7.6, 20 nM DNA. All reactions were quenched by addition of 1M NaOH and 90 °C for five minutes, followed by the addition of loading dye (0.3 M NaOH, 95% formamide, 0.05% bromophenol blue, 0.05% xylene cyanol) and run via 15% PAGE in 1x TBE buffer. Following electrophoresis at 1500 V for approximately 45 min, gels were wrapped and exposed overnight to a storage phosphor screen for image capture. The phosphor screens were scanned using a GE Healthcare Typhoon Trio scanner and quantified via ImageQuaNT (v8.2) software.

DNA binding affinities (K_d_) for MUTYH(Zn-) and MUTYH(Zn+) were determined by electrophoretic mobility shift assays (EMSA) using non-cleavable 2′-fluoro-2′-deoxyadenosine (fA):OG 30-bp duplexes (5 pM final concentration) as previously reported.^17^ For EMSAs, experiments were performed using either *total* enzyme concentration (determined by UV-vis absorbance at 280 nm and extinction coefficient of 84,630 M^-1^) or normalized based on *active* fraction (product of molar concentration and active fraction of enzyme determined by multiple-turnover kinetics experiments). Concentrations of enzyme thus varied from [0-600] nM_280nm_ and [0-300] nM_active_. Binding reactions were prepared using a master mix containing 40 mM Tris–HCl (pH 7.6), 200 mM NaCl, 20% (w/v) glycerol, 0.2 mg/mL BSA, 2 mM DTT, and 20 pM radiolabeled DNA duplex. Equal volumes of the DNA master mix were combined with decreasing concentrations of enzyme, prepared at 4 °C in dilution buffer (20 mM Tris–HCl, pH 7.6, 20% glycerol), and incubated for 30 min at 25 °C. Protein–DNA complexes were separated from free DNA by electrophoresis on 6% native polyacrylamide gels in 0.5× TBE buffer at 120 V for 2 h at 4 °C. Gels were dried, exposed to storage phosphor screens for 48 h, imaged using a GE Healthcare Typhoon Trio scanner, and quantified with ImageQuaNT (v8.2). Binding data were fit by nonlinear regression to a one-site specific binding isotherm to determine the dissociation constant (K_D_), using the equation *Y* = *B*_max_*X*/(*K_D_* + *X*), where *X*is the MUTYH concentration, *Y*is specific binding (%), and *B*_max_is maximal binding.

For comparative analysis, apparent binding affinities (K_1/2_) for all MUTYH proteins were additionally measured by fluorescence polarization using the same reaction mixtures and incubation times and temperatures as EMSA, and with identical duplex DNA except for bearing a 5′-6-FAM label.^57^ Fluorescent DNA (500 pM) was complexed with increasing concentrations of enzyme (1 nM–5 µM), as determined by UV_280nm_ without normalization for active fraction to avoid masking defects in OG recognition, which is required for activity in AST measurements.^17^ Enzyme–DNA complexes were equilibrated for 30 min at 25 °C prior to measurement in 384-well plates using a BMG Labtech Clariostar multimode plate reader (excitation/emission 482/530 nm; gain = 60). All experiments were performed in triplicate, and binding parameters were determined using the one-site specific binding model in GraphPad Prism 10.

### Inductively Coupled Plasma-Mass Spectrometry (ICP-MS) metal analysis

Iron and Zn content of human MUTYH proteins was quantified by inductively coupled plasma–mass spectrometry (ICP–MS). Samples included MUTYH(+Zn), MUTYH(-Zn), H85A, H85C, H85S, C332S, and the R247G/R307G double mutant. Prior to analysis, proteins were buffer-exchanged into 20 mM Tris–HCl (pH 7.5), 250 mM NaCl, and 10% glycerol, and the dialysis buffer was used as a background blank for ICP-MS. Protein samples and buffer blanks were digested in 15% HNO_3_ (v/v) and then diluted with water to a final concentration of 5% (v/v) and heated at 65 °C for 2 h, or until complete digestion was achieved, then filtered through 0.2 µm filters. Resulting digested proteins ranged from 0.4-5 uM. Samples were submitted to the UC Davis Interdisciplinary Center for Inductively Coupled Plasma–Mass Spectrometry for elemental analysis.

### General cell culture

All cell culture experiments were performed using HEK293FT Flp-In cell lines. Wild-type (WT) HEK293FT Flp-In competent cells were provided by the laboratory of Dr. Wolf-Dietrich Heyer (UC Davis). The corresponding MUTYH^-^/^-^ HEK293FT Flp-In cell line was generated in the David laboratory at UC Davis using CRISPR–Cas9 technology, as described previously.^26^ Cells were maintained at 37 °C and 5% CO_2_ in Dulbecco’s modified Eagle’s medium (DMEM; Gibco) containing 4.5 g/L D-glucose and supplemented with 10% fetal bovine serum (FBS; Gibco), 1% non-essential amino acids (Gibco), and 1% GlutaMAX (Gibco).

### Stable cell line generation and validation

Stable cell lines were generated using the Flp-In™ system (Thermo Fisher Scientific). MUTYH variants were introduced into the pcDNA5™/FRT/TO vector by site-directed mutagenesis of the wild-type MUTYH gene. MUTYH expression constructs were co-transfected with the pOG44 Flp recombinase plasmid using Attractene transfection reagent, according to the manufacturer’s instructions. Stable integrants were selected in DMEM supplemented as above and containing 50 µg/mL hygromycin. Media were replaced every 3–4 days following PBS washes until resistant foci appeared (≈2 weeks). Individual colonies were isolated and validated by western blotting **(Figure S4A).**

### Plasmid generation for cellular repair assays

Negative and positive control plasmids (pUC19, GFP OFF, and GFP ON) were propagated in chemically competent DH5α cells and purified using a Qiagen Miniprep kit. The OG:A- reporter plasmid precursor (containing a T:A bp at the lesion position) used to assess MUTYH repair capacity was purified by cesium chloride gradient centrifugation, as previously described.^26^

To generate the OG:A plasmid, 4 µg of CsCl-purified plasmid DNA was nicked with Nb.Bpu10I for 4 h at 37 °C to remove a 29-nt single-stranded fragment, with complete nicking confirmed by 1% agarose gel electrophoresis. An OG-containing oligonucleotide corresponding to the excised sequence (with OG substituted for T; Table 3) was phosphorylated using T4 polynucleotide kinase (New England Biolabs), heat-denatured with the nicked plasmid at 90 °C for 5 min, and slowly annealed to room temperature over ≈2 h. Ligation was performed with T4 DNA ligase (1000 units) for 2 h at room temperature. Residual T:A plasmid was selectively digested using AfeI (25 units) at 37 °C for 16 h, followed by treatment with T5 exonuclease (20 units) for 1.5 h at 37 °C to remove digested and nicked plasmid species. The resulting OG-containing plasmid was purified and concentrated using a NucleoSpin® Gel and PCR Clean-up kit (Macherey-Nagel), according to the manufacturer’s protocol.

### Transfection and flow cytometry

Cells were seeded in 24-well plates at ≈20% confluency 24 h prior to transfection. Each well was transfected with a total of 400 µg plasmid DNA using Attractene transfection reagent, following the manufacturer’s protocol. Plasmids included non-fluorescent pUC19 (negative control), GFP OFF (negative control), OG:A-containing reporter plasmid supplemented with pUC19 carrier DNA (1:3 ratio), or GFP ON (positive control).

Forty-eight hours post-transfection, cells were harvested using 0.05% trypsin-EDTA (Gibco), resuspended in 5% FBS in 1× PBS, filtered through 12 × 75 mm BD Falcon tubes, and maintained on ice prior to analysis. Flow cytometry was performed using a Beckman Coulter CytoFLEX cytometer. Percent OG:A repair was calculated as the fraction of dsRed^+^ cells that were also GFP^+^ and normalized to the GFP ON positive control using the following equation:

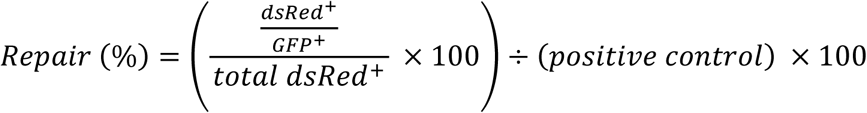

### Multiple sequence and phylogenetic analyses

A multiple sequence alignment including 239 amino acid sequence of Eukaryote MutY homologs and *E.* coli MutY as reference sequence was generated using the MUSCLE algorithm.^58^ The MUSCLE-generated MSA was manually curated and use to construct the phylogenetic tree implementing the neighbor-joining algorithm using 1000 bootstrap replicates. The MSA and the phylogenetic analysis was carried out in the Geneious software package (Version 4.8, Biomatters) and are included in the SI files as ‘MSA.fasta’ and ‘phylogenetic analysis.newick’, respectively. The logo sequence was generated in the WebLogo webpage.

## RESULTS

### Structural overview of MUTYH Zn holoproteins

To provide structural insight into the role of the Zn linchpin motif in OG:A repair, we pursued crystallization trials of Zn-bound MUTYH:DNA complexes reflecting various stages of the proposed adenine excision reaction coordinate. MutY/MUTYH excises adenine by a *retaining* double-displacement mechanism featuring an oxocarbenium-like transition state that is stabilized by transient formation of a covalent glycosyl-enzyme intermediate with the catalytic Asp (Asp 236 in MUTYH). Hydrolysis of the acetal intermediate liberates adenine and generates the β-configured apurinic/apyrimidinic site (AP site).^14,59^ Aza-containing TSAs have been widely used to drive mechanistic and structural studies of MutYs and related glycosylases.^10,14,25,48,59–62^ Among the TSAs, ‘1NBn’ mimics an “early” azaribose transition state and possesses a benzyl moiety appended to a pyrrolidine scaffold, whereas 1-aza-2′-deoxyribose (‘1N’) reflects the cationic oxocarbenium-ion-like transition state formed later in the proposed MUTYH adenine-excision mechanism.^60^ Finally, tetrahydrofuran (THF) is commonly used as a stable mimic of the AP site product formed by monofunctional DNA glycosylases. **Figure 1** summarizes OG:A repair initiated by MutY enzymes, highlighting the proposed mechanism and chemically synthesized mimetics to trap various stages of the reaction coordinate.

**Figure 1.**
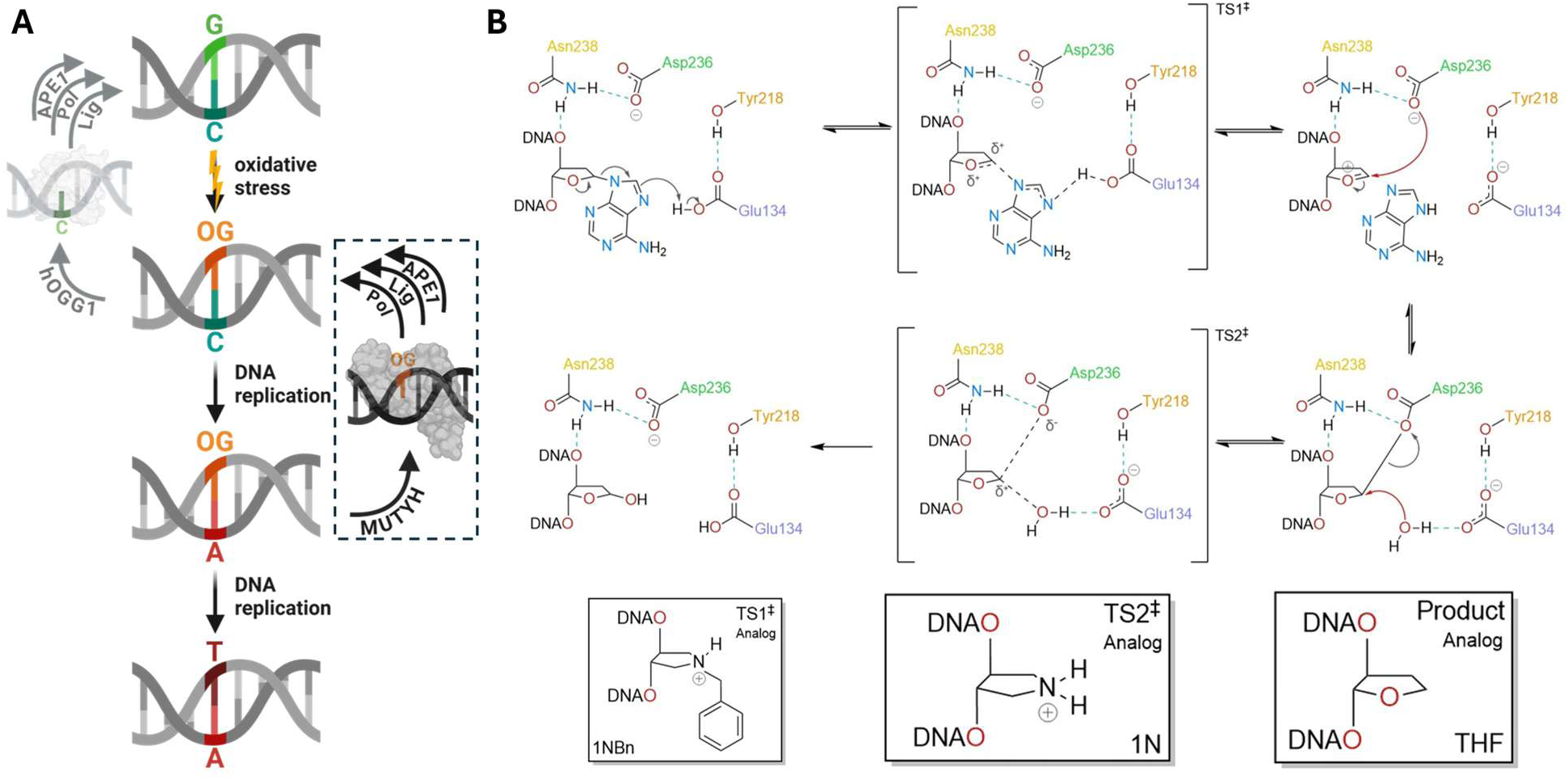
Overview of the guanine oxidation (GO) repair pathway and the proposed MutY adenine excision mechanism. **(A)** GO repair pathway, emphasizing the role of MUTYH in suppressing G:C → T:A transversion mutations by intercepting the post-replicative OG:A lesion and **(B)** the proposed mechanism by which MUTYH cleaves the N-glycosidic bond of mis paired adenines across from OG.

**Figure 1.**
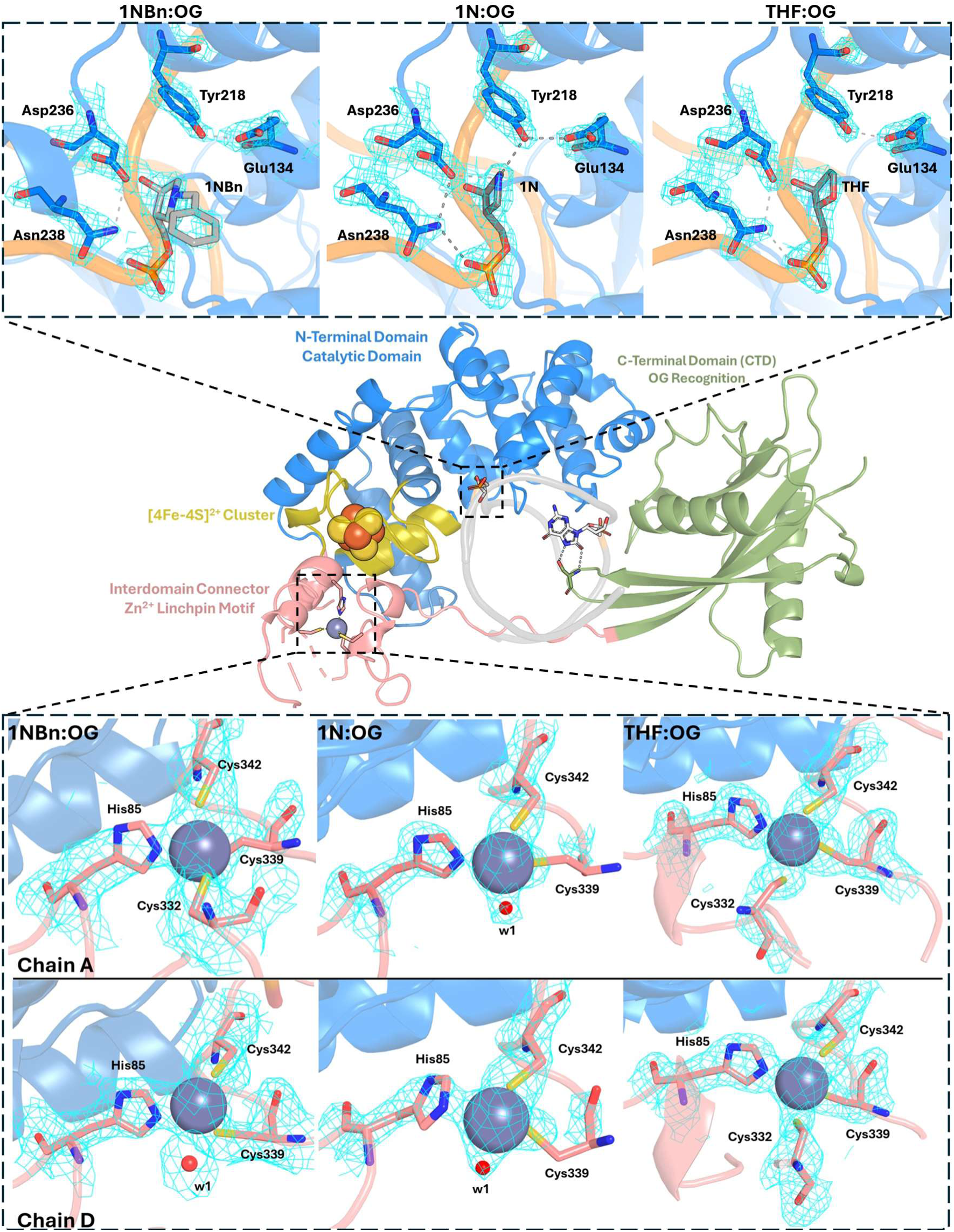
Crystal structures of MUTYH(Zn+) bound to DNA. Structural overview of human MUTYH highlighting its two functional domains: an N-terminal catalytic domain and a C-terminal 8-oxoguanine (OG) recognition domain. A [4Fe-4S] cluster is coordinated within the catalytic domain, while a mononuclear Zn ion resides in the Zinc Linchpin Motif located within the interdomain connector (IDC). **Top:** Simulated annealing omit maps for the active site of MUTYH bound to DNA containing lesions of 1NBn:OG (TSA1), 1N:OG (TSA2), and THF:OG (PAC). **Bottom:** Corresponding simulated annealing omit maps highlighting Zn coordination in MUTYH bound to DNA containing 1NBn:OG (TSA1), 1N:OG (TSA2), and THF:OG (PAC)

Successful X-ray crystallography efforts captured Zn-bound MUTYH proteins complexed with DNA duplexes containing 1NBn and 1N as early and late transition state analogs (TSAs), respectively, or the product analog (PA) tetrahydrofuran (THF), opposite OG as shown by anomalous diffraction scan at the Zn K edge (**Figure S3**). The three structures were determined at a resolution limit ≤ 2 Å by molecular replacement with the MUTYH(Zn-) (Zn-less) structure (PDB ID = 8fay) and refined through iterative cycles of torsion-angle simulated-annealing, restrained minimization, and manual model building. The structures were deposited in the Protein Data Bank with the IDs 11ji (MUTYH-1N:OG), 11jh (MUTYH-1NBn:OG) and 11ip (MUTYH-THF:OG), for the MUTYH containing DNA with 1N, 1NBn and THF, TSA and PA. **Table S1** summarizes the diffraction data processing and refinement statistics. The MUTYH amino acid sequence utilized for crystallization is shown in **Figure S1**, emphasizing residues resolved herein that are not observed in the MUTYH(Zn-) structure.

An overview of MUTYH(Zn+) is shown in **Figure 2**, with enlarged views of the active-site in the top panel and Zn-binding sites from both copies of the MUTYH-DNA complex present in the asymmetric unit (chains A and D). Structural comparison indicates that the benzyl substituent in the 1NBn complex occupies a pre-existing cavity observed in the THF:OG and 1N:OG structures **(Figure S5 left panel)**. To further assess how these analogs engage the active site, we compared their positioning relative to Asp236, the conserved catalytic residue proposed to form a transient covalent intermediate with the C1′ of the deoxyribose **(Figure S5 right panel)**. Overlay of the transition state and product analogs shows that in the transition state analogs (1N/1N-Bn) – where charge buildup is expected at the position corresponding to the anomeric in dA– Asp236 is closely positioned with distances of 2.8 Å for 1N and 3.1 Å for 1NBn. In contrast, this distance increases to 4.2 Å in the product-analog complex. Overall, structural superposition of the active sites across the 1NBn (early TSA), 1N (late TSA), and THF (product-analog) MUTYH-DNA complexes shows that residues adopt nearly identical conformations, consistent with prior structural studies of MutYs.^10,21,25,59^

In the bottom panels of **Figure 2**, the MUTYH-DNA complexes exhibit distinct Zn coordination environments across the different structural states. In the early transition state analog complex (TSAC, MUTYH-1NBn:OG), the coordination sphere is heterogeneous: chain A retains full coordination by all four protein ligands (His85, Cys332, Cys339, and Cys342; HCCC), whereas chain D substitutes Cys332 with a water ligand, hereafter referred to as HOCC. In the late TSAC (MUTYH–1N:OG), which mimics the high-energy oxocarbenium-ion-like transition state that follows base departure, both copies in the asymmetric unit reflect the ‘HOCC’ coordination sphere. By contrast, the product analog complex (PAC; MUTYH–THF:OG), representing the BER intermediate immediately prior to incision by apurinic/apyrimidinic endonuclease 1 (APE1), is the only structure in which both copies in the asymmetric unit retain the complete HCCC coordination sphere, suggesting that the progression of base excision may drive changes in the configuration of the Zn coordination sphere in MUTYH.

### Zn coordination drives distinct conformational states in holo- vs apo- MUTYH

Ligation of the Zn cofactor in MUTYH(Zn+) structures reveals additional organization within the local Zn-binding environment. Zn coordination recruits His85 from the N-terminal domain, resulting in the formation of a new secondary structural element – a 3_10_ helix (H1) encompassing residues Ser82, Ser83, Tyr84, His85, and Leu86 **(Figure S6)**. In parallel, coordination by Cys342 stabilizes surrounding residues within the IDC (aa 317-367). Although Zn binding rigidifies H1 and IDC residues in the immediate coordination sphere, the 8-residue loop connecting Cys332 to Cys339 remain highly disordered in the electron density maps of MUTYH(Zn+).

In all transition state analog complexes (TSAC: MUTYH(Zn+)/1NBn:OG and MUTYH(Zn+)/1N:OG) and product analog complex (PAC: MUTYH(Zn+)/THF:OG), the Zn-binding site of MUTYH is positioned ≈ 16 Å from the [4Fe-4S] cluster and ≈ 24 Å from the active site **(Figure 3A).** In the MUTYH(Zn-) structure, the absence of H1 and disorder within the IDC renders several residues (Val81, Tyr84, Leu86, Phe87, Leu341, Leu343) unavailable for packing against the [4Fe-4S]^2+^ cluster. Instead, the interface between the cluster and the vacant Zn site in MUTYH(Zn-) is partially filled by hydrophobic residues Ala248, Ile249, Ala251, Val258, Leu257, Leu262 and Leu265. Notably, Phe87, which engages in a cation-pi interaction with Arg247 in both Zn- and Zn+ states, undergoes a pronounced χ_2_ rotamer change of ≈90° upon Zn binding, re-orienting itself and the stacked Arg247 residue closer to the [4Fe-4S] cluster. The adjacent Leu86, whose isobutyl side chain points outward (away from the vacant Zn site) in MUTYH(Zn-) rotates to tuck its aliphatic side chain to adopt a more buried conformation in the bound-state **(Figure 3B).** These Zn-dependent rearrangements correlate with a marked reduction in solvent accessibility of the [4Fe-4S] proximal environment in the Zn+ state **(Figure 3C)**. Consistent with this increased exposure in the Zn- state, ICP-MS analysis revealed a reduced Fe/protein ratio of ∼3.3 per monomer in MUTYH(Zn-) compared with ∼4.0 in MUTYH(Zn+), corresponding to an ∼18% decrease in cluster loading in the Zn-deficient protein **(Table 2).**

**Figure 3.**
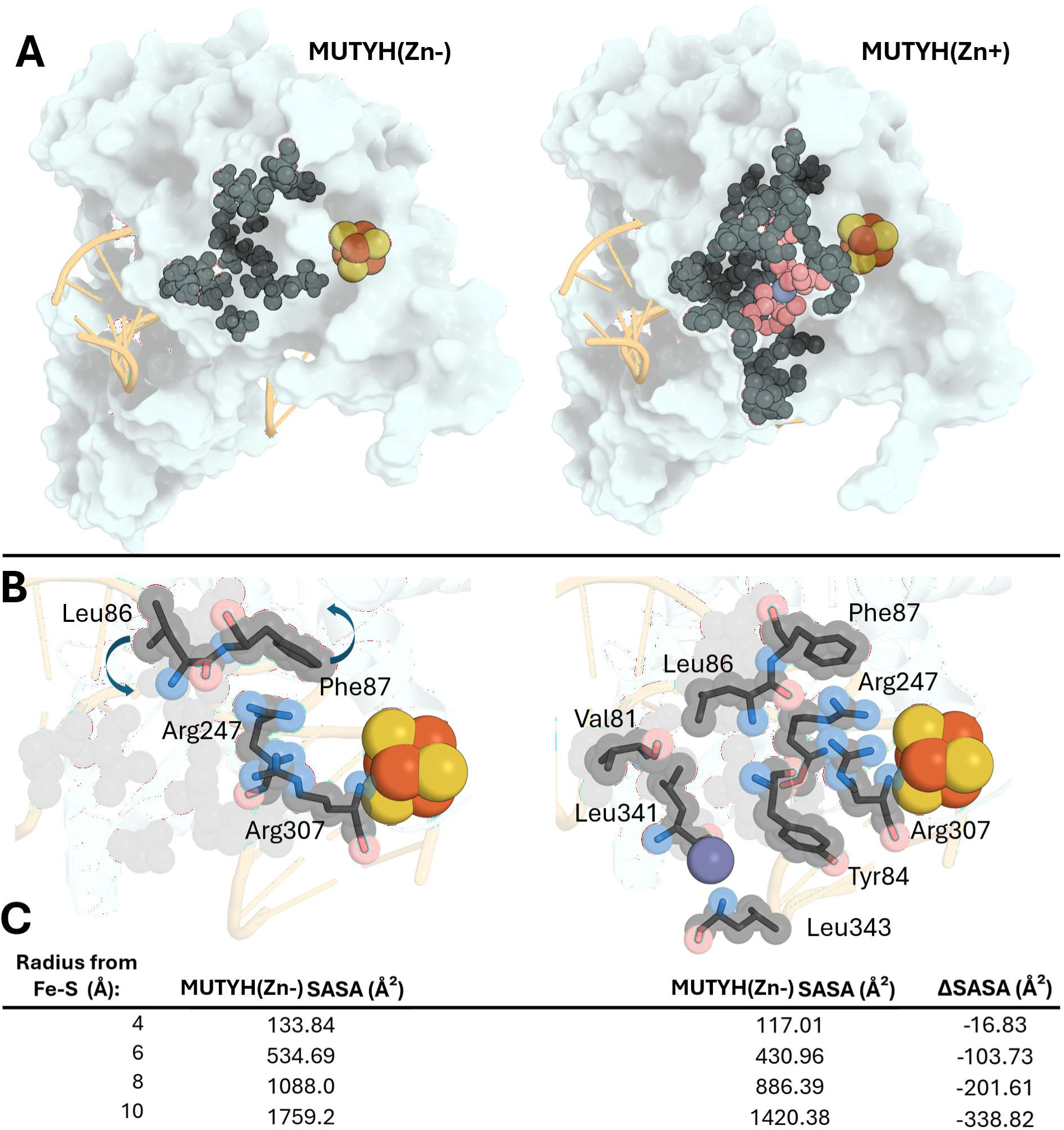
Zn occupancy in MUTYH(Zn+) remodels the solvent-exposed surface surrounding the [4Fe–4S] cluster. **(A)** Residues from the N-terminal domain (H1 and surrounding residues) and the interdomain connector (IDC) ligate and surround the solvent-exposed Zn^2+^ ion (salmon spheres = coordinating residues, grey spheres = non-coordinating residues) in MUTYH(Zn+) models (shown here for the MUTYH(Zn+)–THF:OG complex) **(B)** Sphere & stick model of conformationally dynamic residues in the cofactor interface. The Zn bound state recruits/reorganizes residues Val81, Tyr84, Leu86, Phe87, Arg247, Arg307, Leu341, and Leu343 (dark gray spheres). Ala248, Ile249, Ala251, Val258, Leu257, Leu262 and Leu265 are shown by lighter grey spheres without stick representations, as they are static between the *apo-* and *holo-* models. **(C)** These metalation-dependent changes correspond to the observed reduction in solvent-accessible surface area (SASA) within concentric shells surrounding the [4Fe–4S] cluster as a function of radial distance.

**Table 2.** Summary of *in-vitro* assays.

| Enzyme | <sup>a</sup>K<sub>1/2</sub><br>(nM) | <sup>b</sup>Active<br>(%) | <sup>c</sup>k<sub>2</sub><br>(min<sup>-1</sup>) | <sup>d</sup>k<sub>3</sub><br>(min<sup>-1</sup>) | Zn/<br>protein | Fe/<br>protein |
| --- | --- | --- | --- | --- | --- | --- |
| MUTYH (Zn+) | 17 ± 4 | 51 ± 4 | 1.3 ± 0.2 | 0.001 ± 0.0003 | 1.0 ± 0.1 | 4.0 ± 0.2 |
| MUTYH (Zn-) | 60 ± 8 | 28 ± 5 | 1.1 ± 0.1 | 0.004 ± 0.002 | 0.1 ± 0.1 | 3.3 ± 0.1 |
| H85A | 120 ± 10 | 33 ± 3 | 1.2 ± 0.2 | 0.002 ± 0.001 | 0.5 ± 0.1 | 3.2 ± 0.1 |
| H85C | 86 ± 9 | 35 ± 2 | 1.2 ± 0.1 | 0.003 ± 0.001 | 0.9 ± 0.1 | 3.2 ± 0.1 |
| H85S | 30 ± 3 | 39 ± 2 | 1.3 ± 0.1 | 0.001 ± 0.001 | 0.5 ± 0.1 | 3.8 ± 0.1 |
| C332S | 21 ± 2 | 49 ± 3 | 1.4 ± 0.2 | 0.001 ± 0.0004 | 1.0 ± 0.1 | 3.8 ± 0.1 |
| R247G/R307G | 240 ± 20 | 22 ± 2 | 0.6 ± 0.1 | 0.002 ± 0.001 | 0.4 ± 0.1 | 2.1 ± 0.1 |
| ΔHCCC | 130 ± 12 | 26 ± 3 | 1.2 ± 0.1 | 0.0004 ± 0.0001 | 0.08 ± 0.02 | 3.0 ± 0.2 |

<sup>a</sup>K<sub>1/2</sub> (nM) was determined through fluorescence polarization with incubation temperatures of 25 °C for 30 minutes
<sup>b</sup>active enzyme fractions were extrapolated from the burst amplitude of the respective enzyme's product production curve under multiple turnover conditions at 37 °C
<sup>c</sup>k<sub>2</sub> and <sup>d</sup>k<sub>3</sub> (min<sup>-1</sup>) are derived from the rates of the initial burst phase at 37 °C (under single turnover conditions) and subsequent linear phase (under multiple turnover conditions), respectively.
All values represent means ± S.D of ≥ 3 technical replicates.

A network of polar and covalent interactions connects the Zinc Linchpin Motif, via H1-surrounding residues, to the [4Fe-4S] cluster and active site in MUTYH(Zn+) structures **(Figure 4A)**. His85 donates a hydrogen bond from its ND1 imidazole nitrogen to the backbone carbonyl oxygen of Arg247 (2.7 Å). The backbone carbonyl of the adjacent residue Leu86 forms a polar contact with the guanidinium nitrogen of Arg247, which also engages in a cation-π interaction with the aromatic side chain of Phe87. Arg247 is a terminal residue of catalytic helix H12 and anchors the universally conserved Arg241-Asn238-Asp236 interaction network^10^ to helix H1 and thus, the Zinc Linchpin Motif. In addition, backbone carbonyls of Ser83 and Tyr84 on H1 lie within hydrogen-bonding distance of the Arg307 side-chain nitrogens (NH1/NH2; 3.2 and 2.8 Å, respectively). Arg307, in turn, is immediately C-terminal to the Fe–S cluster–coordinating residue Cys306. Together, Arg247 and Arg307 facilitate contiguous interaction pathways linking the Zn site ∼24 Å to the active site and ∼16 Å to the [4Fe-4S] cluster.

**Figure 4.**
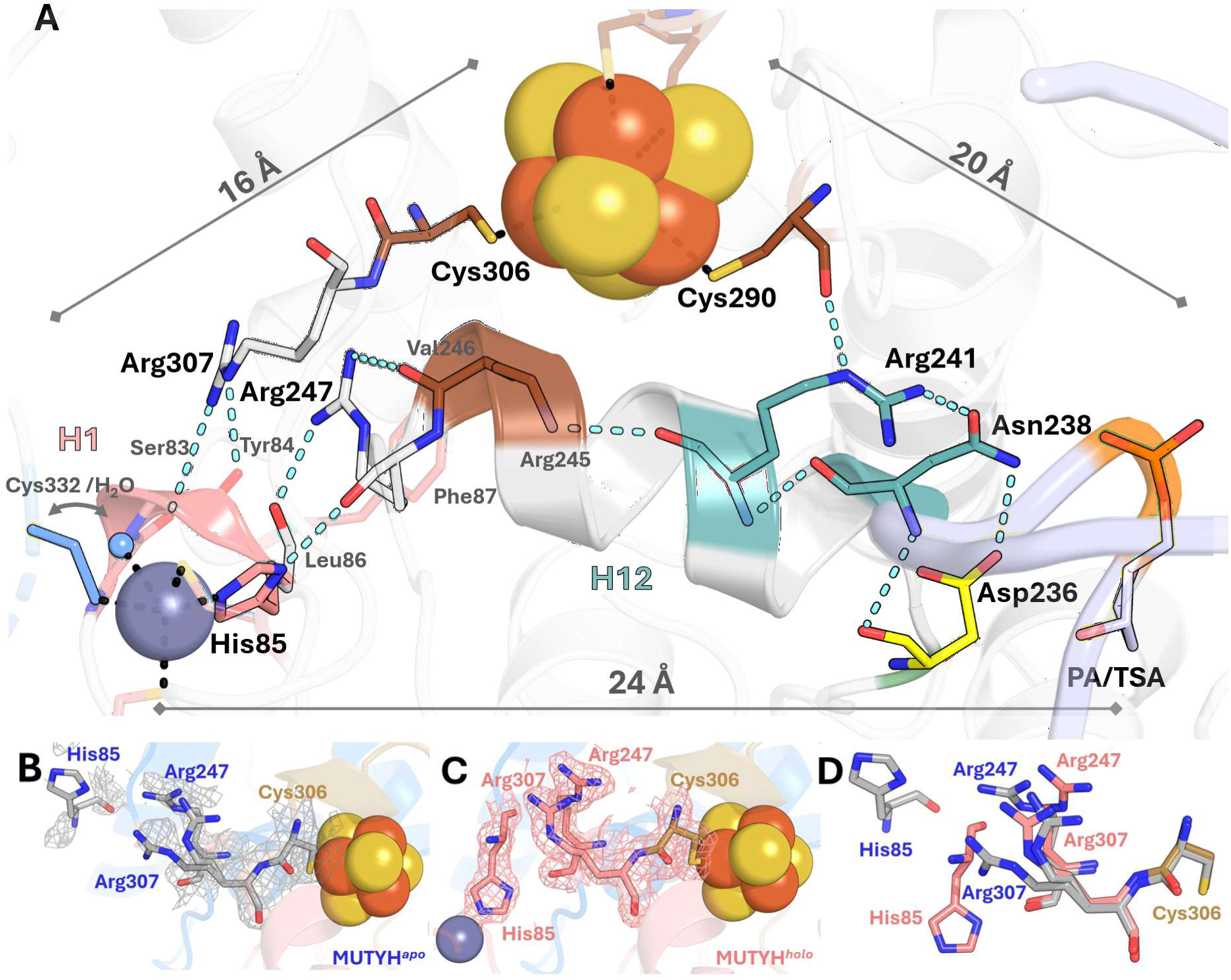
Schematic of the extended network connecting the Zinc Linchpin Motif, [4Fe-4S] cluster, and active site. **(A)** Interconnectivity between the Zinc Linchpin Motif residues and [4Fe-4S] cluster and active site via Arg307 and Arg247 bridging residues, respectively. Simulated-annealing composite omit maps are shown for **(B)** apo and **(C)** Zn-bound MUTYH structures, highlighting differences in local electron density associated with Zn coordination. **(D)** structural overlay of the two preceding panels emphasizes Zn-dependent conformational differences within the His85–Arg247–Arg307 network.

Zn-dependent conformational changes involving helix H1 and the bridging residues Arg247 and Arg307 are visualized in the composite omit maps shown in **Figures 4B-4C**. Loss of Zn leads to disordering of helix H1 and surrounding residues (Ser82–Ser83–Tyr84–His85–Leu86–Phe87; **Figure 4B)**, accompanied by a pronounced displacement of the His85 Cα, which shifts by ∼5.4 Å relative to its position in the MUTYH(Zn+) complexes. The imidazole side chain of His85 is not resolved in the MUTYH(Zn-) structure, consistent with increased local disorder in the absence of Zn. Concomitantly, Arg247 repositions, shifting laterally by ∼3.1 Å toward the vacant Zn site and bending slightly out of plane toward Arg307. The Arg307 side chain becomes highly disordered and was modeled in two alternate conformations: one approximating the holo-like orientation but displaced by ∼1.2 Å away from His85 while remaining approximately coplanar, and a second conformation shifted by ∼2.3 Å toward Arg247, with rotation of the guanidinium group such that one terminal nitrogen (NH1/NH2) points toward Arg247 while the opposing terminal nitrogen is oriented downward. Collectively, Zn coordination is associated with a defined conformational arrangement of Arg247 and Arg307 that accommodates helix H1 residues within the Zinc Linchpin Motif **(Figures 4C**, **4D).**

### Zn linchpin or Arg247/Arg307 perturbation impairs MUTYH activity and cellular repair

To benchmark the Zn-deficient enzyme against the holoprotein, expression and purification conditions were optimized (detailed in Methods section) to modulate Zn incorporation, yielding MUTYH(Zn+) holoprotein as well as MUTYH(Zn-) preparations. Metal occupancy of purified proteins was quantified using inductively coupled mass spectrometry (ICP-MS, results summarized in **Table 2**).

We first compared DNA binding by electrophoretic mobility-shift assay (EMSA; **Figure S7**) of MUTYH(Zn-) and MUTYH(Zn+) preparations under low DNA conditions to estimate the intrinsic dissociation constant (K_d_) for Zn-dependent binding to 5’ γ-^32^P labeled, OG:fA lesion containing DNA. Subsequently, fluorescence polarization on 5’ 6-carboxyfluorescein (6-FAM) labeled OG:fA containing DNA was used to determine relative binding affinities (K_1/2_) of unnatural MUTYH variants as a comparative measure to probe Zn binding and bridging sites. Binding experiments were carried out either using absolute enzyme concentration, determined by its molar concentration approximated by UV-Vis spectroscopy at 280nm, or the enzyme concentration normalized to the active population of enzyme determined by the amplitude of the enzymatic burst phase observed under multiple-turnover conditions. This provides a practical estimate of the ‘active fraction’ of catalytically competent enzymes and therefore serves as a benchmark of enzyme behavior prior to slow steady-state turnover, where subtle defects in DNA binding or lesion engagement can be manifested as changes in the population of productive enzyme-substrate complexes rather than alterations in intrinsic catalytic and turnover rate constants. When analyzed by total enzyme concentration (based on UV absorbance at 280nm), the apparent equilibrium dissociation constant (K_d_) for MUTYH(Zn-) on OG:fA (a non-cleavable OG:A analog) DNA substrate was 12 ± 2 nM, approximately three-fold weaker than that of MUTYH(Zn+) (4 ± 0.4 nM). When normalized to equal concentrations based on enzyme active fraction the difference was largely eliminated (0.6 ± 0.1 nM vs 0.8 ± 0.09 nM). This convergence indicates that the apparent binding defect of MUTYH(Zn-) arises from a reduced population of DNA-engaged enzyme molecules, rather than from intrinsic and irreconcilable defects in DNA binding or OG:A lesion recognition (such as mutations of catalytic or OG-recognition residues). Thus, loss of Zn decreases the fraction of enzyme capable of productive lesion engagement without measurably altering binding by the competent population.

To extend these observations beyond forcible Zn removal, we disrupted the Zinc Linchpin Motif through mutations of His85 and bridging residues Arg247 & Arg307. Replacement of His85 to Ala, Cys, or Ser produced distinct biochemical and metal-loading phenotypes **(Table 2**; **Figure 5B).** All three variants exhibited reduced active-enzyme fractions (A_0_) relative to WT MUTYH(Zn+) (51 ± 4%), with H85A being most compromised (33 ± 3%) and H85C and H85S displaying modestly higher activity (35 ± 2% and 39 ± 2%, respectively). Despite these differences in active fraction, all variants exhibited comparable single-turnover rates (*k*_2_ ≈ 1.1–1.3 min^-1^) and indistinguishable steady-state turnover rates (*k*_3_ ≈ 0.001–0.003 min^-1^). Notably, only the serine substitution (H85S) substantially rescued DNA ^OG:fA^ binding affinity (K_1/2_ = 30 ± 3 nM), approaching that of WT MUTYH(Zn+) (17 ± 4 nM) (Figure 5A). In contrast, H85C and H85A exhibited significantly weakened binding (86 ± 9 nM and 120 ± 10 nM, respectively). Inductively coupled plasma–mass spectrometry (ICP-MS; **Table 2**) revealed strikingly different metal-loading profiles: H85A was depleted of both cofactors (0.5 Zn / 3.2 Fe), H85C largely retained Zn but lost Fe (0.9 Zn / 3.2 Fe), whereas H85S preserved Fe but was Zn-deficient (0.5 Zn / 3.8 Fe).

**Figure 5:**
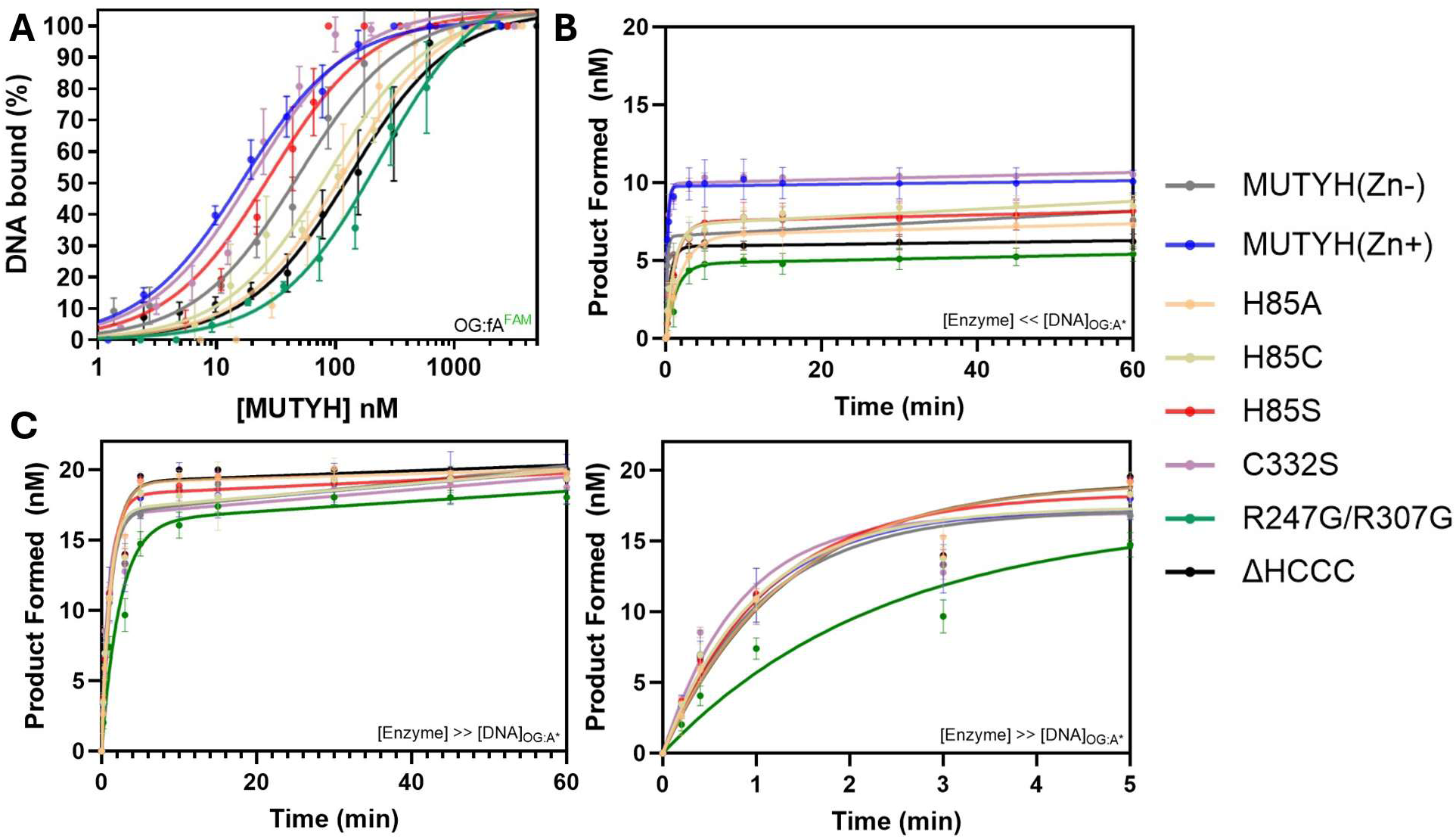
In vitro binding, activity, and kinetic assays performed on MUTYH proteins. **(A)** Fluorescence polarization (fP) was utilized using a range of enzyme concentrations (determined by UV_280nm_) against 500pM of OG:fA (a non-cleavable OG:A analogue) to determine the relative binding affinities (K_1/2_) for MUTYH proteins. **(B)** Gel-based glycosylase assays were performed with excess OG:A DNA substrate to access burst amplitude (relating to active population of enzyme) and the slow turnover step, k_3_ (min^-1^) and **(C)** with excess enzyme (under single-turnover conditions) to ascertain burst kinetics and the rate of catalysis, k_2_ (min^-1^). An expanded view of the first 5 minutes of the single-turnover assays is shown to the right of **(C).**

To evaluate the cellular repair capacity of the H85A, H85C, and H85S variants, corresponding MUTYH^-^/^-^ HEK293FT Flp-In cell lines were generated as described in the Methods. Cells were transfected with a GFP-based reporter plasmid containing a site-specific OG:A lesion within the GFP coding sequence, such that fluorescence depends on MUTYH-initiated repair and completion of downstream base excision repair. A schematic of the assay workflow is shown in **Figure 6A**, with representative flow cytometry plots in **Figure 6B**. Overall, cellular repair efficiencies **(Figure 6C)** were consistent with in vitro data underscoring the indispensability of His85, with H85A exhibiting the most severe defect (56% repair), followed by H85S (59%) and H85C (69%), the latter showing partial but incomplete recovery relative to WT MUTYH (93%).

**Figure 6.**
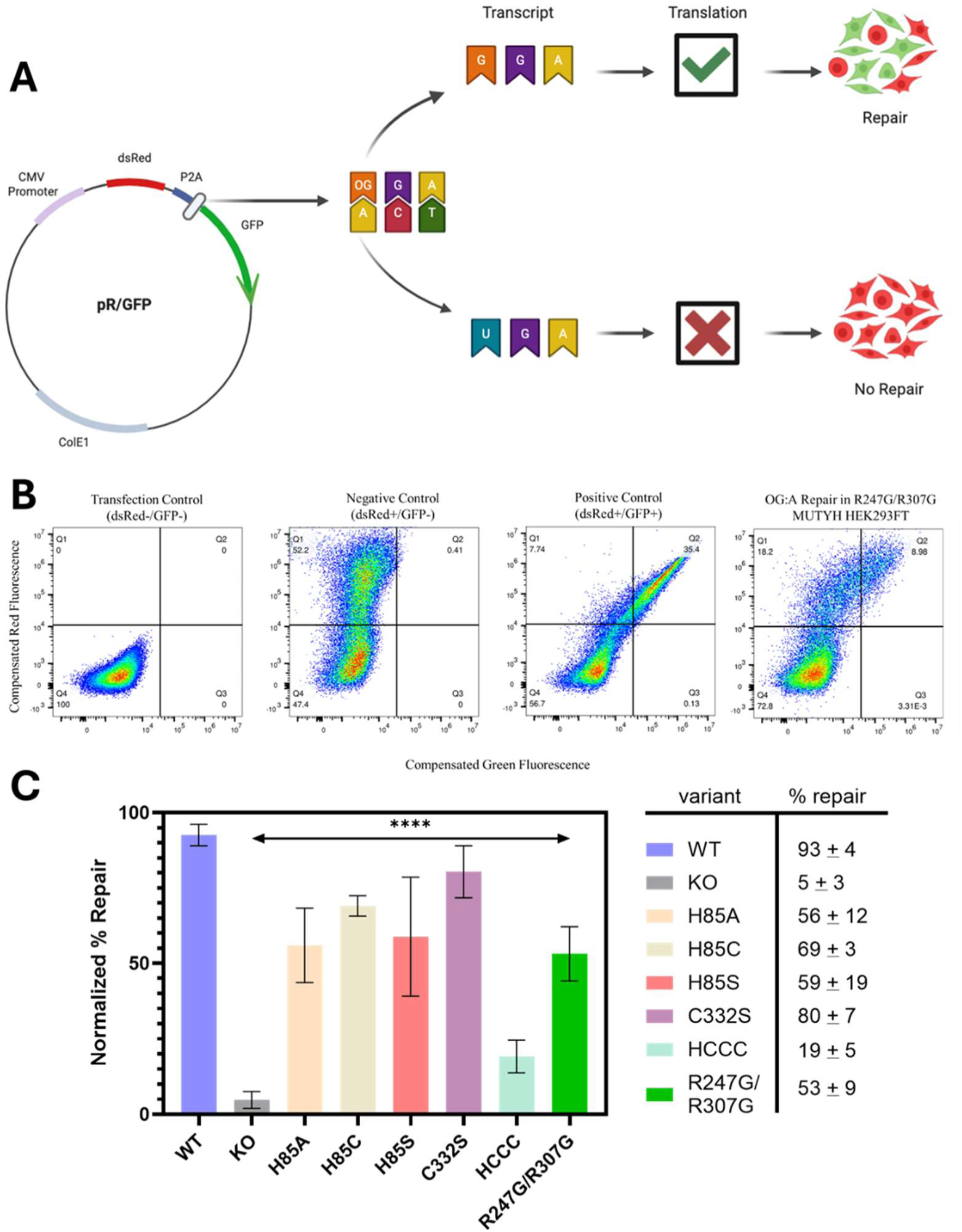
Cellular OG:A repair assay in HEK293FT cell lines expressing MUTYH variants. **(A)** Schematic of the GFP-based OG:A reporter plasmid. Transfected cells constitutively express dsRed, while successful repair of an OG:A lesion within the downstream GFP gene by MUTYH and subsequent base excision repair (BER) is required to permit GFP expression. **(B)** Representative flow cytometry plots of compensated dsRed (y-axis) versus GFP (x-axis) fluorescence in HEK293FT cells quantifying MUTYH-mediated OG:A repair. From left to right, plots show the transfection control pUC19 (empty vector; dsRed^-^/GFP^-^), the negative control containing a T:A base pair at the lesion position (dsRed⁺/GFP⁻), the positive control containing a G:C base pair at the lesion position (dsRed^+^/GFP^+^), and a representative MUTYH R247G/R307G variant. Quadrants correspond to untransfected cells (lower left), transfected dsRed^+^ cells (upper left), and repair-positive dsRed^+^/GFP^+^ cells (upper right); GFP-only cells were not observed. **(C)** Quantification of repair capacity in MUTYH variant HEK293FT cell lines. Data are presented as mean repair efficiency normalized to the positive control (%) ± S.D. Statistical significance relative to wild-type MUTYH was assessed using one-way ordinary ANOVA, with **** indicating *p* < 0.0001.

To assess the consequences of complete Zn loss in cells, we additionally generated a quadruple mutant (H85A/C332S/C339S/C342S; hereafter ΔHCCC) designed to abolish Zn coordination in cells. Strikingly, the ΔHCCC variant exhibited a dramatic loss of cellular OG:A repair (19 ± 5%), far exceeding the modest reductions observed for individual Zn-site mutants and contrasting sharply with the relatively mild effects on glycosylase activity observed with recombinantly expressed and purified ΔHCCC, whose *in vitro* profiles resembled those of MUTYH(Zn-) and individual His85 mutants **(Figure 5**, **Table 2).** Repair by cells expressing the MUTYH ΔHCCC mutant establishes a functional benchmark for Zn deficiency in cells, demonstrating that Zn loss in cells is substantially more detrimental in OG:A repair than simply the more modest effects the metal cofactor plays in OG:A recognition and adenine excision *in vitro,* suggesting a role for the metal cofactor in downstream steps of BER.

Arg247 and Arg307 sit at the interface between the Zn-binding site, the catalytic core, and the [4Fe–4S] cluster. Arg247 is positioned at the Zn-proximal end of helix H12, which encompasses the Arg241–Asn238–Asp236 active-site network^10^ **(Figure 4A)**, while Arg307 extends toward the [4Fe–4S] cluster. To assess their functional importance, we generated the R247G/R307G double-substitution to maximally increase local flexibility and disrupt both backbone- and side-chain–mediated connectivity across both paths of the interface. Although Arg247 and Arg307 lie outside the primary coordination spheres of both metal cofactors, the double mutant exhibited pronounced loss of both Zn and Fe (0.4 Zn and 2.1 Fe per protein). Among all variants tested, R247G/R307G exhibited the poorest *in vitro* biochemical profile. The substrate analog OG:fA duplex affinity was markedly reduced (K_1/2_ = 240 ± 20 nM) which is reflected in the low active fraction (A_0_ = 22 ± 2%) measured under multiple-turnover conditions. Remarkably, even under DNA-saturating conditions ([MUTYH]_active_ ≫ [DNA]_OG:A_), where other Zn-perturbed variants exhibited near-wild-type *k*_2_ values, the catalytic rate of R247G/R307G was impaired by ∼2-fold (*k*_2_= 0.6 ± 0.1 min^-1^; Figure 4C, Table 2). Consistent with its in vitro profile, R247G/R307G exhibited markedly reduced cellular repair capacity (53 ± 9%; **Figure 6C).**

### Arg247/Arg307 bridge evolved in concert with the emergence of the Zn-Linchpin Motif

To evaluate whether Arg247 and Arg307 are evolutionarily associated with the Zn-binding site, we performed a phylogenetic and sequence analysis of eukaryotic MutY homologs using a rooted phylogenetic tree and corresponding amino acid sequence alignment. All sequences analyzed retained the conserved Fe–S cluster motif **(Figure 7).** Previous work from our group and others extensively characterized the structural and biochemical features of the Zinc Linchpin Motif in mammalian MutY homologs, particularly in mouse Mutyh.^30,32,33^ Extending this analysis across eukaryotes revealed that conservation of the Zn coordination sphere (H/C/C/C) extends beyond mammals to other vertebrates, including reptiles (birds, turtles, and snakes) and aquatic vertebrates (bony fish, amphibians, and cartilaginous fish). Notably, in all vertebrate MutY clades that retain the H/C/C/C binding motif, Arg247 and Arg307 are largely conserved, with the exception of some rodent homologs in which Arg307 is substituted by Glu. In contrast, many non-vertebrate eukaryotic MutY homologs (e.g., slime molds, arthropods, and stramenopiles) exhibit only partial conservation of the H/C/C/C motif, often with variation at the first Cys ligand (H/X/C/C). In these groups, the residues bridging the [4Fe–4S] cluster and Zn site are also only partially conserved; for example, Arg247 is retained in arthropods, whereas Arg307 is frequently substituted by His or Ser. By contrast, MutY homologs from plants, green algae, nematodes, and several protozoan lineages show little to no conservation of either the Zn-coordinating ligands or the Arg247/Arg307 pair, consistent with loss of the Zn-binding site in these clades. In Archamoebae, only the second and third Cys ligands are retained (X/X/C/C), suggesting that these residues represent vestigial remnants of a degenerated Zinc Linchpin Motif.

**Figure 7.**
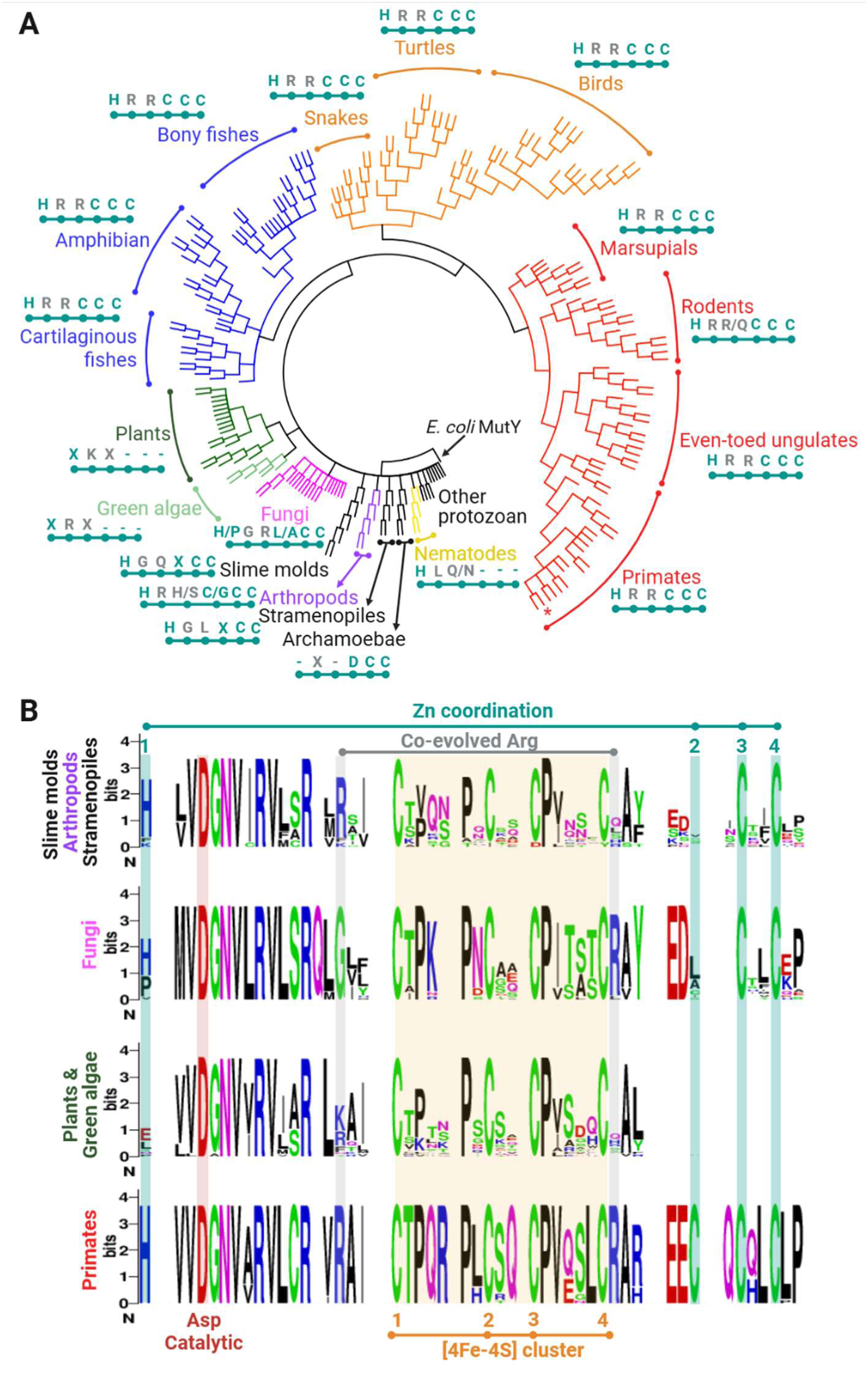
Evolutionary analysis of the Zn coordination motif including Arg247 and Arg307 in eukaryotic MUTYH homologs. **(A)** Phylogenetic analysis of Fe–S cluster–containing MutY homologs reveals that the emergence of the second metal-binding motif (H/C/C/C) coincides with the conservation of Arg247 and Arg307. This suggests an evolutionary linkage between acquisition of the Zn site and the retention of these arginine residues, consistent with a role in structurally coupling the Zn site to the [4Fe-4S] cluster. **(B)** Sequence logo analyses from representative taxa (arthropods/slime molds/stramenopiles, fungi, plants/green algae, and primates) illustrate that conservation of the [bridging arginine residues is restricted to primates, which – among these taxa shown – uniquely possess the Zn coordination motif.

### Dynamic Zinc Coordination Organizes a Protein-Protein Interaction Surface on the IDC

Given the structural differences in Zn coordination across the reaction-coordinate mimics (early TSAC [1NBN:OG], late TSAC [1N:OG], and PAC[THF:OG]) and prior screens of Cys332 mutations in mouse Mutyh,^12,33^ we surmised that the transiently coordinating Cys332 is dispensable for glycosylase activity and OG:A repair. We assessed serine replacement (C332S) at this position and found no measurable change in Zn or Fe content by ICP-MS **(Table 2)**, consistent with the lack of coordination of Cys332 to the Zn site in MUTYH TSAC models. Furthermore, in vitro functional profiling of C332S resembled WT MUTYH(Zn+) behavior for: DNA-binding (K_1/2_ = 21 ± 2 nM vs 17 ± 4 nM in WT), glycosylase active fraction (A_0_ = 49 ± 3 % vs 51 ± 4% in WT) and kinetic parameters (k_2_ ≈ 1.4 vs 1.3 min^-1^ in WT, k_3_ ≈ 0.001 for both enzymes). Notably, while glycosylase activity and metal retention were unchanged *in vitro*, cellular OG:A repair – which reflects the operation of the full base excision repair pathway – was modestly reduced for C332S (80 ± 7% vs 93 ± 4% repair for WT in HEK293FT cells; **Figure 6C).** Of note, C332R is a MUTYH CAV that we have similarly found resembles WT in terms of in vitro activity yet exhibits even more dramatically diminished OG:A repair in cells (64 ± 14%) than C332S.^63^ Together, these results indicate that Cys332, while not required for Zn coordination or intrinsic glycosylase activity, may contribute to overall OG:A repair efficiency in the cellular BER context.

Crystallographic B-factor analysis indicates that Zn binding partially stabilizes the interdomain connector (IDC) relative to the apoprotein, with residues in and around the Zn Linchpin Motif exhibiting lower overall mobility in MUTYH(Zn+) than in MUTYH(Zn-) **(Figure S8, Table S2).** In particular, residues 80–90 encompassing Zn coordinating residue His85 and the short 3_10_ helical component, the cluster-bridging residues Arg247 and Arg307, and segments of the IDC (residues 320–350) display reduced B-factors in Zn+ states, consistent with local rigidification proximal to the Zn site near the surface of MUTYH. Notably, this stabilization is not uniform: the Cys332-X_6_-Cys339 loop retains elevated B-factors across all structures, indicating persistent local disorder despite overall Zn-dependent rigidification of the surrounding IDC scaffold. In concordance with Zn loss being associated with a less ‘relaxed’ state, far-UV circular dichroism measurements (200–240 nm; **Figure S9A**) reveal reduced mean residue ellipticity and broader spectral features for MUTYH(Zn-) relative to MUTYH(Zn+), consistent with increased local conformational heterogeneity in the absence of Zn. In parallel, differential scanning fluorimetry **(Figure S9B)** shows that retention of Zn increases the thermal stability of MUTYH, shifting the melting temperature (T_m_) by ∼3.1 °C.

Given the solvent-exposed positioning of the Zn ion in the flexible IDC, the observed ligand exchange between Cys332 and water in the product analog complex (PAC), persistent disorder within the Cys332-X_6_-Cys339 loop (where X_6_ represents the six disordered amino acids between the Cys332 and Cys339 ligands), and prior reports implicating the IDC as a protein-protein interaction platform for APE1,^27–29^ we explored whether this Zn-dependent surface could participate in MUTYH-APE1 association. To this end, we used AlphaFold3^53,64^ Multimer to generate structural predictions of the MUTYH-APE1-DNA ternary complex. The AlphaFold models were filtered according to the conformation of active site residues, DNA engagement, and Zn coordination **(Figure S10A-C)**. Notably, AlphaFold currently does not model complex cofactors such as the [4Fe-4S] cluster, however, it renders a faithful structural approximation of the [4Fe-4Fe] cluster motif and orientation of its cysteinyl ligands **(Figure S10B)**. Given the intrinsic association of both MUTYH and APE1 towards DNA, the length of the DNA sequence (containing a central THF:OG lesion) was elected to be 90 bps to provide enough DNA-protein interaction to avoid forcible MUTYH-APE1 binding.

The ternary complex shows MUTYH and APE1 adjacently positioned and entirely bound to DNA **(Figure 8)**. Particularly, MUTYH occupies the THF:OG lesion, flipping the THF out into the active site **(Figure S10A)** as observed in the MUTYH-THF:OG crystal structure. This suggests that despite the presence of APE1, MUTYH shields the AP site for further processing by APE1. Interestingly, the IDC region that is rigidified by Zn coordination (residues 320–350), is presented as the main MUTYH-APE1 surface interaction, contributing several polar contacts via backbone amides Ala319 and Ser320, and side chain ofAsp350 (**Figure 8**). Additionally, MUTYH’s PIP box – a short motif that mediates binding to the proliferating cell nuclear antigen (PCNA) – mapped at the end of the C-terminal domain, docks (Leu529, Phe532 and Phe533) into a hydrophobic pocket of APE1 flanked by hydrogen bonds donated by Arg523 and His536 from the C-terminal domain of MUTYH. The involvement of the IDC in the AlphaFold model in the interaction with APE1 is in accordance with biochemical and structural data derived from this work and others,^27–29^ while the implication of the PIP box has not been previously reported. Previous studies have, however, described MUTYH’s PIP box as a region that interacts with PCNA’s IDCL motif.^65,66^ Thus, these computational studies suggest that the PIP-box might mediate multiple protein-protein interactions in MUTYH. Taken all together, our studies suggest that the interactions with the PIP box of MUTYH along with interaction mediated by distinct configurations of the Zn coordination may regulate MUTYH interaction with APE1, likely during handoff of the AP site intermediate.^63^ Indeed, this idea is consistent with our proposal that compromised hand-off to APE1 is the origin of reduced cellular OG:A repair of several MUTYH CAVs within the Zn linchpin/IDC (including C332R) despite WT-like in vitro activity.

**Figure 8.**
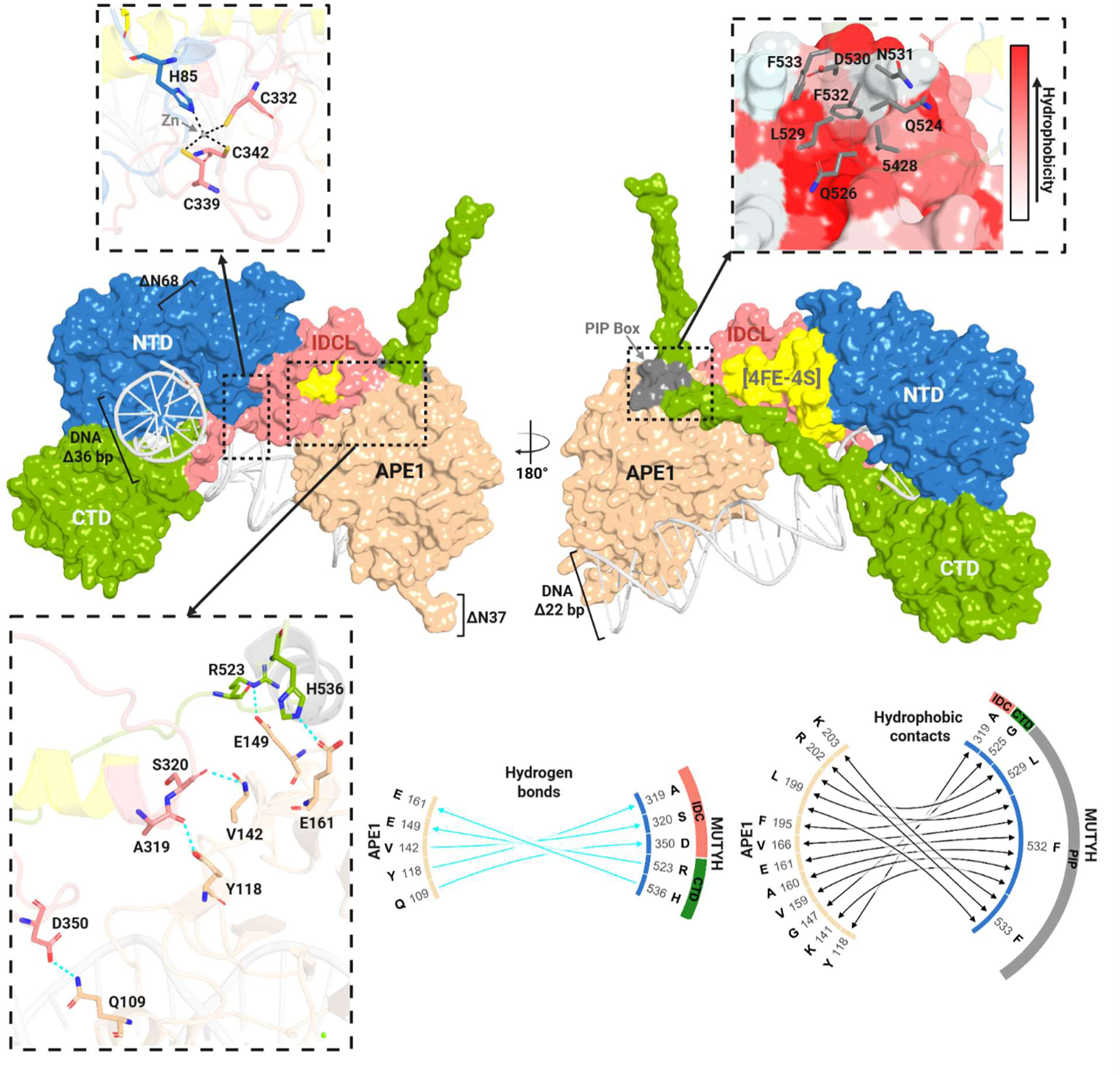
AlphaFold3 model of the ternary MUTYH-APE1-DNA complex. MUTYH domains are depicted in distinct colors: blue, N-terminal domain (NTD); C-terminal domain (CTD), green; Interdomain connector loop, light red; [4Fe-4S] cluster motif, yellow; PIP box, gray. APE1 is shown in wheat color. The Zn coordination sphere and predicted important regions for MUTYH and APE1 interaction are highlighted. MUTYH’s PIP box interaction is shown in terms of hydrophobic interactions, where APE1 region involved in such interaction is colored based on hydrophobicity. The circus plots showcase the hydrogen and hydrophobic interactions involved in MUTYH-APE interaction. The hydrogen bonds are shown in cyan.

## DISCUSSION

The three full-length human MUTYH-DNA structures reported in this work capture distinct catalytic states while retaining both the [4Fe-4S] and Zn cofactors **(Figure 2)**. Across complexes that mimic sequential stages of the reaction coordinate (early-TSAC, late-TSAC, PAC), the active-site residues superpose closely, with analogue-dependent shifts in geometry relative to the catalytic residue Asp236. This behavior parallels prior MutY-family structural snapshots spanning lesion recognition and transition state mimicry, which likewise emphasize a largely conserved active-site scaffold across the reaction coordinate.^21,59^ Notably, in the MUTYH-1NBn:OG complex, the benzyl substituent is accommodated in a pre-existing cavity (“hole”) within the active site without requiring reorganization of surrounding residues, indicating that this added steric bulk can be tolerated within the active-site architecture **(Figure S5 left panel).** In our structures, the distance between Asp236 and the anomeric-mimetic nitrogen is 3.1 Å for 1NBn and 2.8 Å for 1N, whereas this separation increases to 4.2 Å in the THF-bound (product-analog) complex **(Figure S5 right panel)**. The shorter Asp–1N distance is consistent with the 2.9 Å contact reported for *Geobacillus stearothermophilus* MutY (GsMutY) bound to DNA containing 1N:OG.^59^ In prior analyses, 1N is interpreted as an aza-sugar transition state analog that when protonated mimics development of oxocarbenium-like character at the anomeric center, and close approach of the catalytic Asp is proposed to maximize electrostatic stabilization in this state. Within this framework, the slightly longer Asp–1NBn distance in our complex is compatible with a more substrate-like configuration relative to 1N, while remaining within hydrogen-bonding/ion-pairing distance of Asp236.

Comparison of the Zn-bound and Zn(-) structures illustrates the role of Zn coordination in organizing the structure of the interdomain connector (IDC) adjacent to the [4Fe-4S] cluster. Relative to the MUTYH(Zn-) model, the Zn-bound structures preserve helix H1 and stabilize a majority of the IDC, enabling residues to pack against the [4Fe-4S] cluster shielding it from solvent and stapling the [4Fe-4S] cluster domain to the DNA substrate **(Figure 3B)**. Conversely, IDC disorder in the absence of Zn renders these residues unavailable for cluster-adjacent packing, providing a structural rationale for why Zn stripping (e.g., EDTA treatment) yields an [4Fe-4S] cofactor that is more solvent exposed **(Figure 3C)** and partially Fe-depleted **(Table 2).** Such exposure may labilize and degrade the [4Fe-4S] cluster as has been seen in other metalloenzymes.^67–70^ Thus, the presence of Zn provides direct and indirect influence DNA binding of the [4Fe-4S] cluster via domain packing and shielding the [4Fe-4S] cluster.

A central insight from these findings is that the Zn linchpin is structurally coupled to both the [4Fe-4S] cluster DNA binding domain and the catalytic residues that mediate base excision through a long-range interaction network. His85, together with the conserved Arg247 and Arg307 pair, links the Zn site (and H1) to helix H12, which encompasses the Arg241-Asn238-Asp236 active-site network^10^, and to the [4Fe-4S] cluster environment adjacent to Cys306 (**Figure 4A)**. This contiguous pathway provides a structural basis for long-range communication between metal cofactors and the active site, consistent with prior functional studies demonstrating that perturbations in the [4Fe-4S] cluster environment can influence DNA binding and lesion recognition without directly altering the chemistry of adenine excision. ^15–20,34–40^ Specifically, His85 mutagenesis studies suggested that simply substituting for another metal coordinating residue, in the case of H85C, did not completely restore glycosylase activity, metal loading or cellular OG:A repair. Taken together, these data indicate that His85 plays dual roles in MUTYH function, as has been observed for histidine ligands in other metalloproteins, ^71–75^ serving both as a Zn coordinating ligand and as a structural element within an inter-cofactor network that spans to the active site, tuning efficient DNA repair. Furthermore, the severe functional consequences of simultaneously disrupting both Arg247 and Arg307 underscore the importance of Zn linchpin motif integration with both the active site and the [4Fe–4S] cluster DNA binding domain.

The Zn coordination sphere itself exhibits state-dependent changes across the reaction coordinate, with ligand exchange at Cys332 distinguishing transition state-like and product-like complexes. The WT-like profile of C332S *in vitro* **(Table 2)** supports prior biochemical and mutational work showing that Zn coordination by Cys332 is comparatively dispensable for glycosylase activity.^12,33^ The absence of Cys332 coordination in the late TSAC, which captures MUTYH in a catalytically committed post-base departure state, followed by its re-engagement in both copies of the PAC, suggests that clamping of the Zn coordination sphere by Cys332 is temporally coupled to product formation and may poise the IDC for downstream handoff events. This is evidenced further by the reduced cellular OG:A repair associated with mutation of this ligand to Ser **(Figure 6C)** and Arg^76^, together with the pronounced OG:A repair defects observed by the removal of all Zn coordinating ligands (ΔHCCC, **Figure 6C**).^63^ Thus, the pronounced repair outcomes observed with Zn loss in cells, in contrast to inconsequential (i.e. C332S/R) or modest (ΔHCCC) glycosylase behavior *in vitro* (**Figure 5**) indicate an additional role of the Zn linchpin motif in MUTYH in downstream steps of base excision repair beyond OG:A lesion detection and adenine base removal.

Consistent with this view, Zn binding partially rigidifies the IDC scaffold while preserving localized flexibility within the Cys332-X_6_-Cys339 loop. This dynamic architecture is well suited for regulated protein-protein interactions and aligns with prior reports implicating the IDC as a platform for coordinating downstream BER factors, including APE1, the 9-1-1 clamp, and SIRT6.^27–30^ AlphaFold prediction of the ternary MUTYH-APE1-DNA complex suggests that APE1 engages the Zn-containing face of the IDC, converging on a surface shaped by Zn-dependent organization yet retaining flexibility at the ligand-bearing loop. This duality, coupled with active-site dependent dynamics at the Zn coordination sphere / APE1 binding interface, provides a plausible structural mechanism by which Zn coordination and active site product formation could pre-organize MUTYH for productive handoff to downstream repair enzymes while accommodating conformational heterogeneity required for transient interactions.

Our structural, biochemical, and cellular data support a model in which His85 serves as a key organizing element of the Zn → active site and the Zn→ [4Fe-4S] communication network via Arg247 and Arg307, while Cys332 functions as an active-site-dependent exchangeable Zn ligand that is dispensable for *in vitro* glycosylase activity but influences cellular repair outcomes by tuning the conformational properties of the IDC. Recent large-scale saturation mutagenesis of MUTYH analysis of OG:A repair activity in cells largely agrees with our results herein in showing compromised repair upon alteration of the Zn coordinating ligands, though less deleterious to repair than mutations of the [4Fe-4S] cluster Cys ligands.^12^ **Figure 9** summarizes that, together, these observations position the Zn linchpin as an allosteric organizer that integrates [4Fe–4S] cluster integrity and DNA binding, active-site network connectivity, and protein-protein interaction capacity, thereby coordinating long-range structural interplay required for efficient MUTYH-mediated DNA repair in the cellular environment.

**Figure 9.**
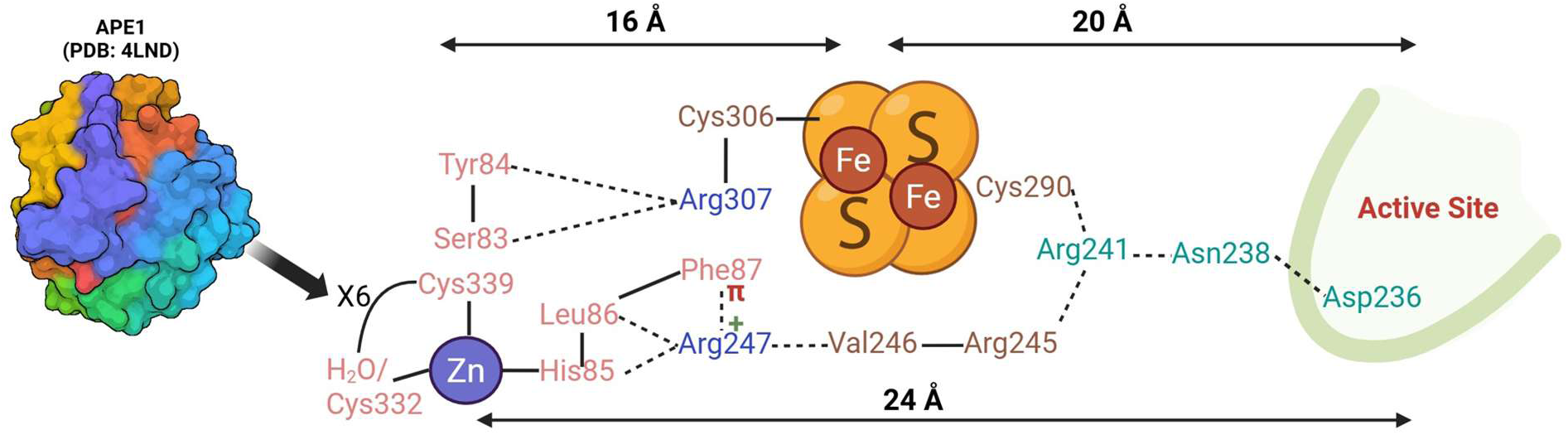
Schematic of the extended network connecting the Zn Linchpin Motif, [4Fe-4S] cluster, and active site. A simplified 2D schematic depicts the network of interactions extending approximately 16 Å from the Zn binding site to the [4Fe–4S] cluster, and a further 20 Å from the cluster to the catalytic active site, illustrating long-range connectivity between the two metal cofactors and the site of catalysis. Numerous covalent (solid lines) and polar contacts (dashed lines) from H1 in the N-terminal domain (residues 83-87, salmon) mediate interactions with bridging residues Arg307 and Arg247 which span 16 Å to the cluster and 24 Å to the active site, respectively. A highly disordered loop between the transient ligand Cys332 spans six residues to coordinating residue Cys339, which is hypothesized to mediate interactions with APE1 for product hand-off.

From an evolutionary standpoint, the co-emergence of the Zn linchpin motif with the Arg247/Arg307 bridging network in vertebrate MUTYH homologs supports the notion that this inter-cofactor communication pathway represents a regulatory adaptation layered onto the conserved MutY scaffold.^30,32,33^ In contrast, non-vertebrate eukaryotic homologs that lack full conservation of the Zn ligands also show reduced conservation and dependence on bridging residues Arg247 and Arg307 **(Figure 7)**. The ability to fine-tune MUTYH activity in humans and other animals is likely necessary to precisely control adenine excision and coordinate AP site handoff, limiting the accumulation of these genotoxic abasic site intermediates that are prone to strand breaks and drive genomic instability in multicellular organisms. As such, the risk of systemic consequences such as cancer may have imposed evolutionary pressure for the expansion of the conserved MutY scaffold described herein. Indeed, MUTYH has been suggested to play dual roles in preventing carcinogenesis by thwarting accumulation of DNA mutations and initiating cell death pathways via strand breaks under in precancerous cells under conditions of high oxidative stress. ^77–79^ Regulation and accuracy in fulfilling these multiple roles thereby relies on the Zn linchpin’s dual roles in facilitating lesion engagement and regulating hand-off of the AP site to APE1. Moreover, the reliance on these functions in preventing carcinogenesis is further underscored by CAVs that disrupt the Zn-linchpin network, such as those at His85, Arg247 and Arg307, and compromise repair.

## Supporting information

MSA.fasta

phylogenetic analysis.newick

## Acknowledgements

We thank the Interdisciplinary Center for Plasma Mass Spectrometry at UC Davis for providing access to ICP-MS instrumentation used for metal quantification in this study. In particular, we thank Austin Cole for technical assistance, detailed analysis, and mentorship in operating the instrument during the course of this work.

## Author contributions

MH: Conceptualization [equal], Data curation [lead], Formal analysis [lead], Writing – original draft [lead], Writing – review & editing [equal], MM: (Data curation [equal], Writing – review & editing [equal] Formal analysis [supporting], Writing – review & editing [supporting]), TX: (Data curation [equal]), AL: (Data curation [supporting]), AXLC: (Data curation [supporting]), AF: (Formal analysis [supporting], Supervision [supporting], Validation [equal]), CHTA: (Conceptualization [equal], Data curation [equal], Formal analysis [equal], Project administration [equal], Supervision [equal], Writing – original draft [equal], Writing – review & editing [equal]), and SSD (Conceptualization [equal], Formal analysis [equal], Project administration [lead], Supervision [lead], Writing – original draft [equal], Writing – review & editing [lead])

## Data availability

Coordinates and structure factors have been deposited in the Protein Data Bank, https://www.rcsb.org/ (PDB IDs: 11ji, 11jh and 11ip)

