## supplemental information for "Structural Connectivity Between the Zinc Linchpin Motif, the [4Fe-4S] cluster, and the active site orchestrates DNA repair in MUTYH"

* Corresponding author:

Sheila S. David. University of California, Davis, One Shields Avenue, Davis, CA 95616, U.S.A.

.

Proposed journal: Nucleic Acids Research

Key words: MutY, Base Excision Repair, DNA glycosylase, [4Fe-4S] cluster, Zinc Linchpin Motif, allostery.

**TABLE OF CONTENTS**

**Supplementary Figures**

**Figure S1. Protein sequence of the MUTYH construct used for crystallographic studies.**

**Figure S2: UV–visible absorption spectra of MBP–MUTYH fusion proteins expressed under different zinc supplementation regimes.**

**Figure S3: Representative X-ray absorption scan collected from MUTYH(Zn+) crystals in complex with THF:OG containing duplex DNA.**

**Figure S4. Western blot analysis of HEK293FT cell lines and recombinant MUTYH proteins generated in this work.**

**Figure S5. Transition-state analog binding preserves the MUTYH active-site architecture.**

**Figure S6. Two-dimensional secondary structure representation of human MUTYH(Zn+) in complex with THF:OG-containing duplex DNA, generated using PDBsum**

**Figure S7. Zinc-dependent DNA binding affinities of MUTYH measured by electrophoretic mobility shift assay (EMSA).**

**Figure S8. Residue mobility across MUTYH structures**

Supplementary information

**Figure S9. Effects of Zn Binding on MUTYH Thermal Stability and Secondary Structure**

**Figure S10. Structural comparison of the functional component between AlphaFold model and crystallography MUTYH structures.**

### <u>Supplementary Tables</u>

**Table S1: Crystallographic Statistics**

**Table S2** Composite-region mean B-factors in MUTYH(Zn+) and MUTYH(Zn-)

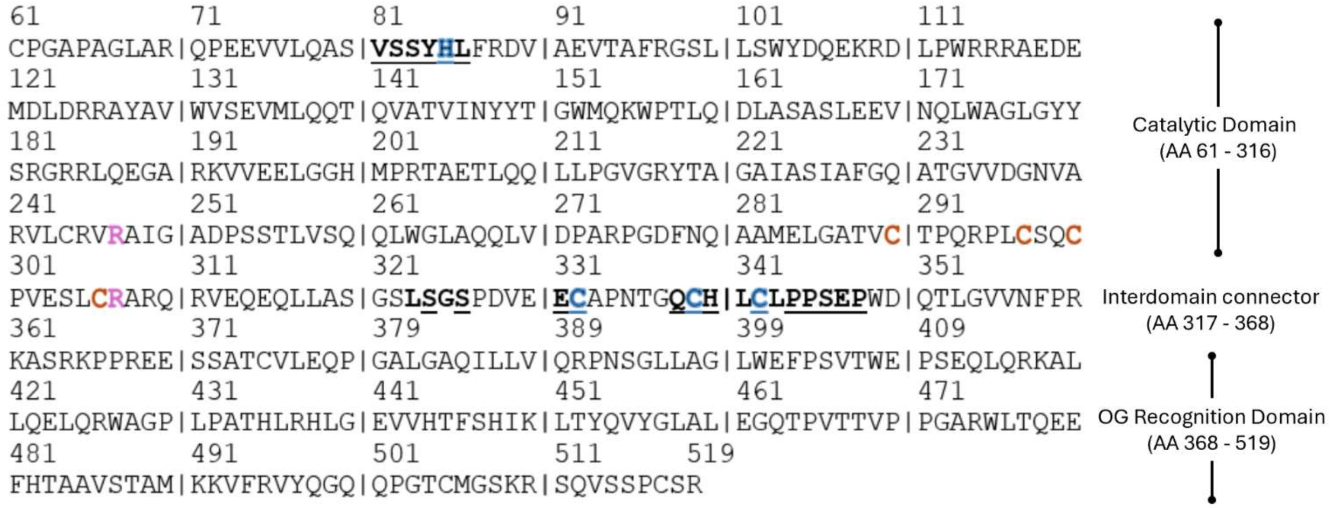


**Figure S1. Protein sequence of the MUTYH construct used for crystallographic studies.** Residues rendered in **bold** correspond to regions resolved in the crystal structures presented in this study, which were not resolved in existing MUTYH models; but specific residue resolution may vary depending on the bound DNA lesion or transition-state analog (e.g., THF versus TSA complexes). **Underlined** residues denote loci associated with cancer-linked variants. Residues highlighted in **blue** indicate Zn-coordinating ligands, **brown** indicate [4Fe–4S] cluster ligands, and **pink** identify arginine residues that bridge the zinc and iron–sulfur cofactors.

Supplementary information

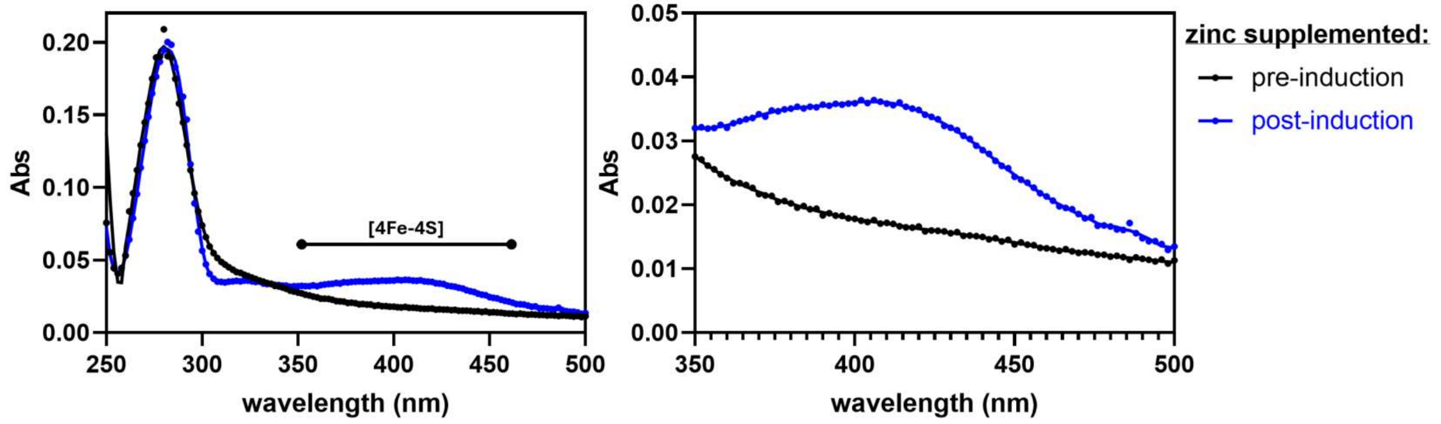


**Figure S2: UV visible absorption spectra of MBP–MUTYH fusion proteins expressed under different zinc supplementation regimes.** Proteins were expressed in *E. coli* with 200uM (final concentration) of $ZnSO_4$ supplementation either immediately prior to IPTG induction (black trace) or 20 h post-induction (blue trace). **Right**, magnified view of the 350–500 nm window, highlighting the broad absorption centered at ~410 nm that is characteristic of the [4Fe–4S] cluster in MUTYH.

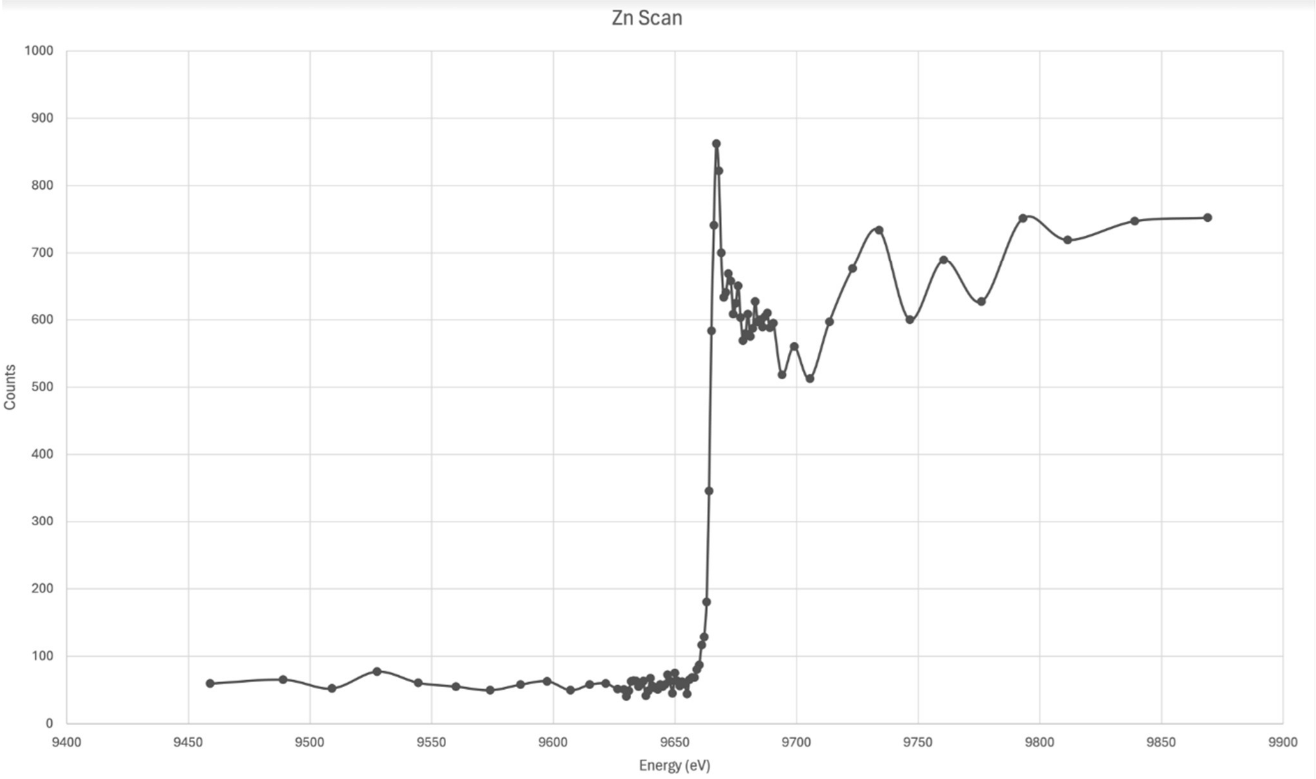


**Figure S3: Representative X-ray absorption scan collected from MUTYH(Zn+) crystals in complex with THF:OG–containing duplex DNA.** The signal shows a clear rise and maximum near the Zn K-edge (~9660 eV), confirming the presence of zinc in the MUTYH(Zn+) DNA complex

Supplementary information

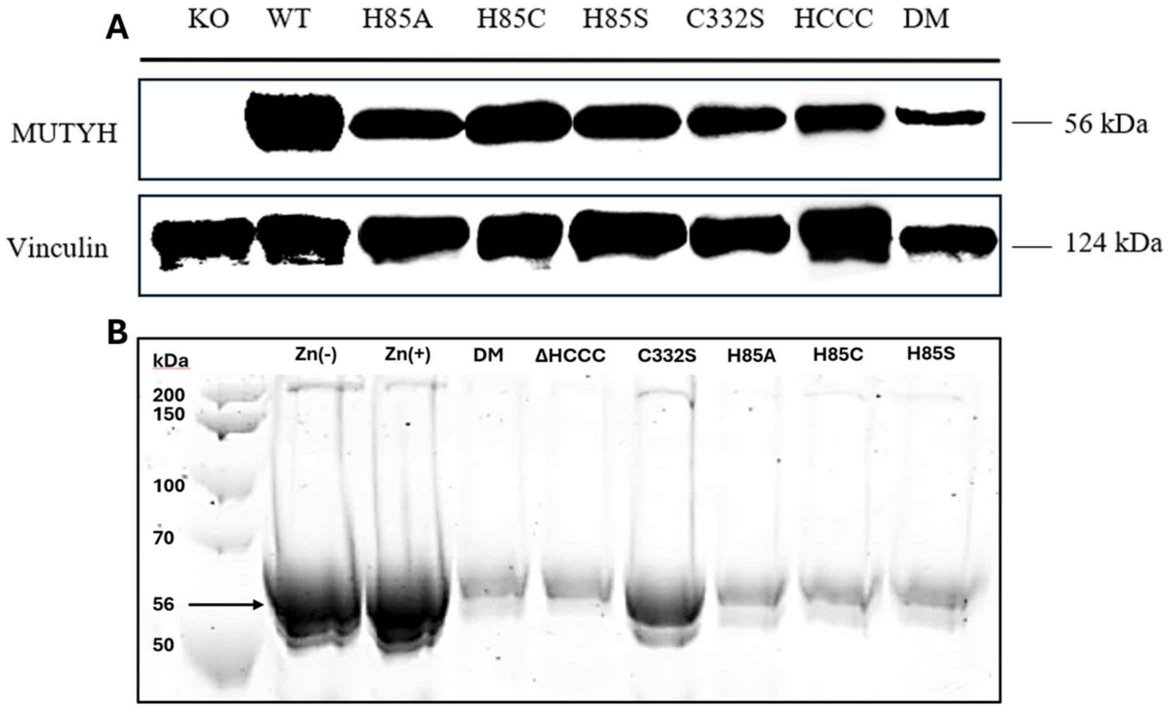


**Figure S4. Western blot analysis of HEK293FT cell lines and recombinant MUTYH proteins generated in this work. (A)** From left to right: $MUTYH^{-/-}$ knockout HEK293FT cell line, WT HEK293FT, H85A, H85C, H85S, C332S, H85A/C332S/C339S/C342S quadruple mutant (ΔHCCC), R247G/R307G double mutant. **(B)** From left to right: WT MUTYH(Zn-), WT MUTYH (Zn+), DM, ΔHCCC, C332S, H85A, H85C, H85S

Supplementary information

| Table S1: Crystallographic Statistics | | | |
|---|---|---|---|
| **Structure** | **MUTYH-1N:OG** | **MUTYH-1NBn:OG** | **MUTYH:THF:OG** |
| **PDB ID** | **11ji** | **11jh** | **11ip** |
| Beamline | SSRL 12-1 | SSRL 12-1 | SSRL 12-1 |
| Wavelength | 1.00003 Å | 0.97946 Å | 0.979460 Å |
| Resolution range (Å) | 38.87 - 1.85 (1.88 - 1.85) | 37.62 - 2.00 (2.04 - 2.00) | 38.76 - 1.70 (1.73 - 1.70) |
| Space group | $P\ 2_1\ 2_1\ 2_1$ | $P\ 2_1\ 2_1\ 2_1$ | $P\ 2_1\ 2_1\ 2_1$ |
| Unit cell | *a* = 87.95Å, *b* = 116.76Å, *c* = 118.22Å | *a* = 88.01Å, *b* = 117.41Å, *c* = 118.02Å | *a* = 87.63Å, *b* = 116.69Å, *c* = 118.30Å |
| Total reflections | 359,747 (18,027) | 421,826 (24,640) | 798,830 (36,673) |
| Unique reflections | 99,551 (5,013) | 79,814 (4,583) | 127,779 (5,789) |
| Multiplicity | 3.6 (3.6) | 5.3 (5.4) | 6.3 (6.3) |
| Completeness (%) | 96.1 (98.3) | 96.3 (97.5) | 96.0 (89.0) |
| Mean I/sigma(I) | 7.9 (1.3) | 8.4 (1.9) | 14.0 (1.6) |
| Wilson B-factor | 29.65 $Å^2$ | 36.14 $Å^2$ | 24.88 $Å^2$ |
| R-merge | 0.073 (1.125) | 0.099 (0.776) | 0.067 (1.102) |
| R-meas | 0.085 (1.305) | 0.115 (0.878) | 0.073 (1.199) |
| R-pim | 0.042 (0.641) | 0.047 (0.365) | 0.029 (0.467) |
| CC1/2 | 0.997 (0.632) | 0.995 (715) | 0.999 (0.616) |
| Reflections used in refinement | 99,454 (10,119) | 79,746 (7,994) | 127,709 (12,077) |
| Reflections used for R-free | 4,938 (516) | 3,935 (408) | 6,344 (589) |
| R-work | 0.1703 (0.2997) | 0.2966 (0.3850) | 0.1683 (0.2554) |
| R-free | 0.2116 (0.3521) | 0.3384 (0.4112) | 0.2000 (0.2769) |
| Number of non-hydrogen atoms | 8,036 | 7,467 | 8,247 |
| macromolecules | 7,248 | 7,147 | 7,356 |
| ligands | 94 | 115 | 71 |
| solvent | 718 | 253 | 844 |
| Protein residues | 813 | 816 | 824 |
| RMS(bonds) | 0.011 Å | 0.023 Å | 0.011 Å |
| RMS(angles) | 1.12° | 1.91° | 1.21° |
| Ramachandran favored (%) | 97.88 | 94.26 | 97.77 |

| | | | |
|---|---|---|---|
| Ramachandran allowed (%) | 1.99 | 4.86 | 2.10 |
| Ramachandran outliers (%) | 0.12 | 0.87 | 0.12 |
| Rotamer outliers (%) | 0.45 | 2.46 | 0.74 |
| Clashscore | 5.19 | 16.26 | 3.71 |
| Average B-factor | 39.06 Å$^2$ | 44.13 Å$^2$ | 33.33 Å$^2$ |
| macromolecules | 37.29 Å$^2$ | 42.77 Å$^2$ | 31.58 Å$^2$ |
| ligands | 51.21 Å$^2$ | 40.50 Å$^2$ | 46.07 Å$^2$ |
| solvent | 43.75 Å$^2$ | 39.56 Å$^2$ | 39.55 Å$^2$ |

Statistics for the highest-resolution shell are shown in parentheses.

Supplementary information

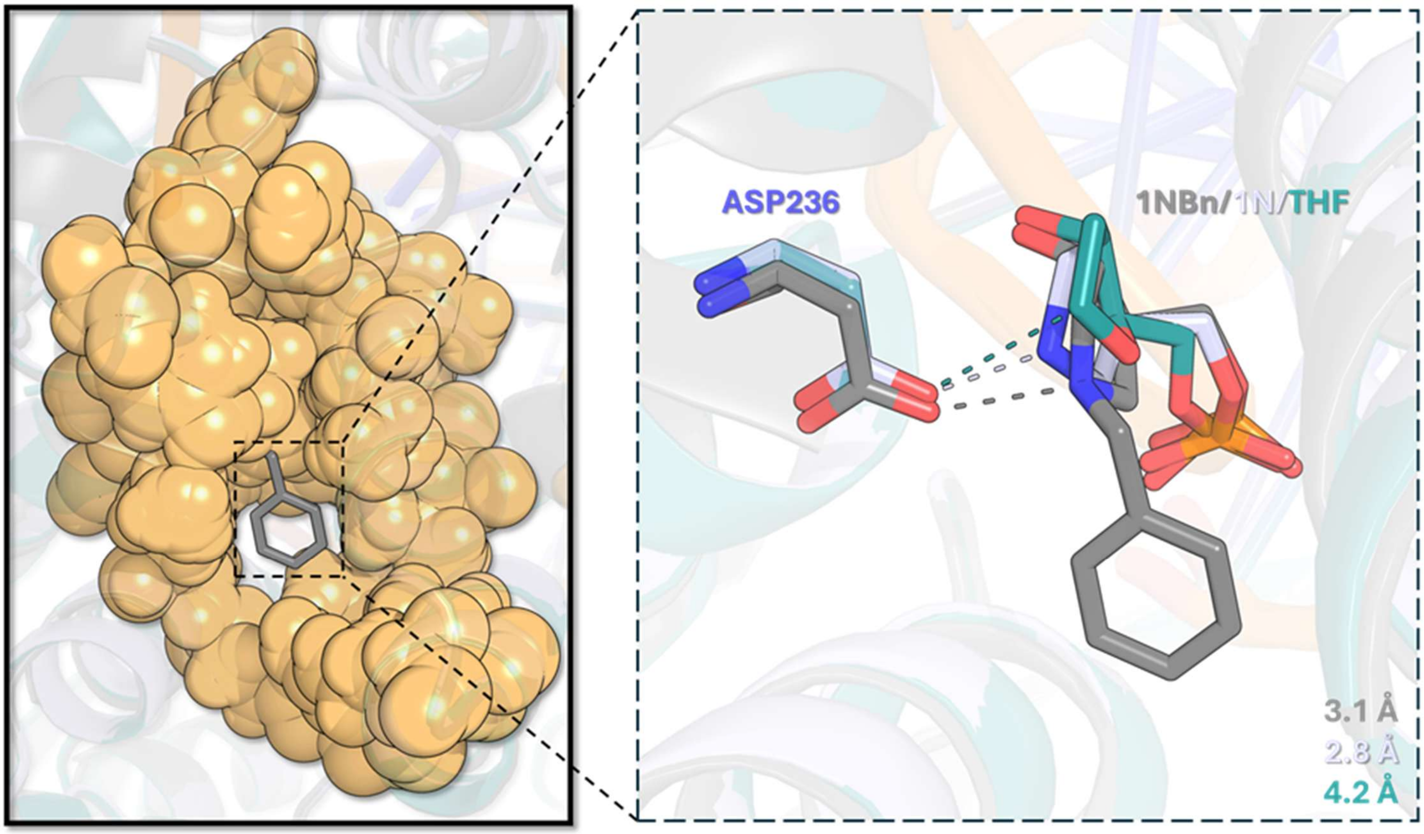


**Figure S5. Transition-state analog binding preserves the MUTYH active-site architecture. Left**, comparison of MUTYH structures bound to different analogs reveals that the TS1 analog 1NBn occupies a pre-existing cavity within the active site without inducing detectable conformational rearrangements. In contrast, the baseless sugar and product analog complexes (1N:OG and THF:OG, shown as orange space-filling models) exhibit a void within this region, which is filled by the benzyl substituent in the 1NBn structure. Superposition of active-site residues across all complexes indicates no measurable changes in side-chain conformations (not shown). **Right**, overlay of the analogs used in this study highlighting distances between the catalytic nucleophile Asp236 and the anomeric center of the sugar. In the transition-state analog complexes, which mimic charge buildup at the anomeric position, Asp236 is positioned in close proximity (2.8 Å for 1N and 3.1 Å for 1NBn), whereas this distance increases to 4.2 Å in the product analog (THF) complex, consistent with loss of covalent interaction following catalysis.

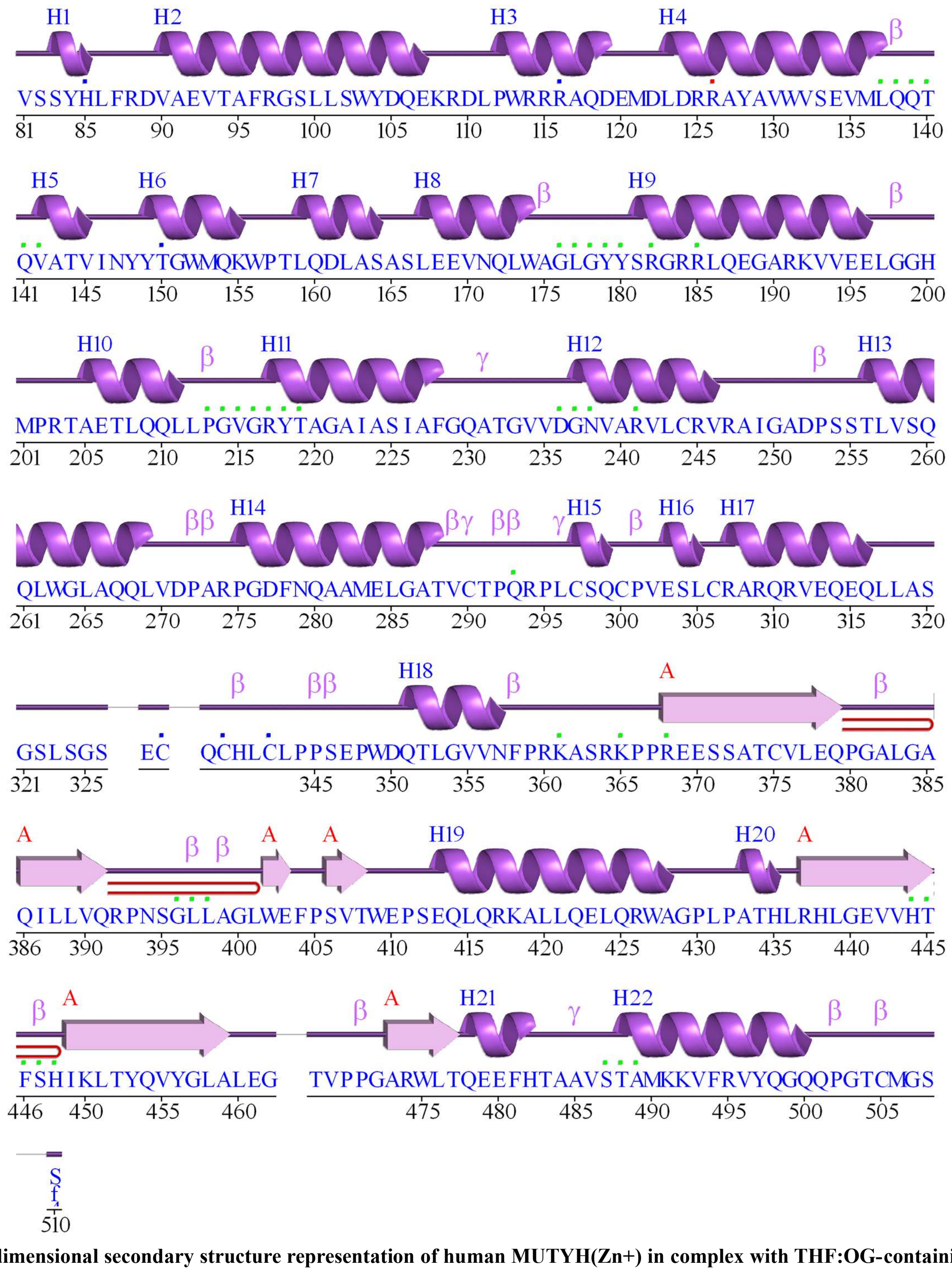


**Figure S6**. **Two-dimensional secondary structure representation of human MUTYH(Zn+) in complex with THF:OG-containing duplex DNA, generated using PDBsum.** α-Helices (H1-H22) are shown as purple cylinders, β-strands are indicated by arrows, and connecting loops are shown as lines. Residue numbering is shown below the sequence. Regions corresponding to β-hairpins and extended strands are annotated, highlighting the predominantly α-helical architecture of MUTYH interspersed with short β-elements. The distribution of secondary structural elements illustrates the modular organization of the protein scaffold that supports lesion recognition and catalytic activity in the Zn+ complex.

A)

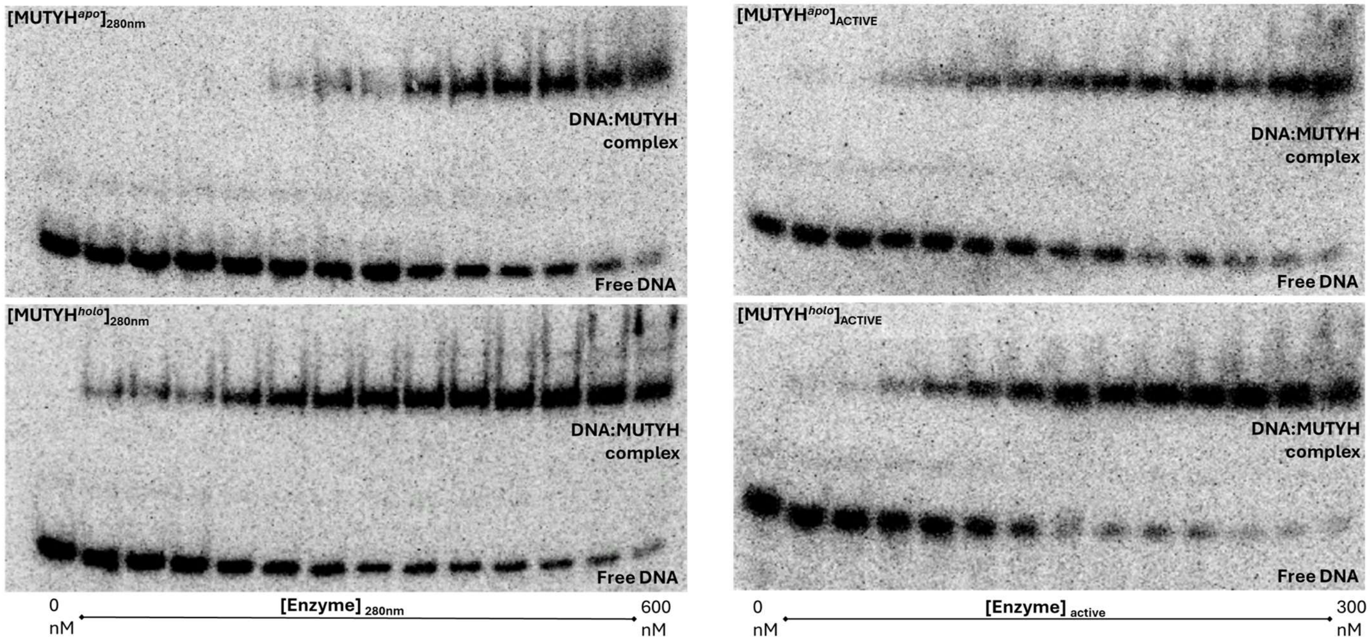


B)

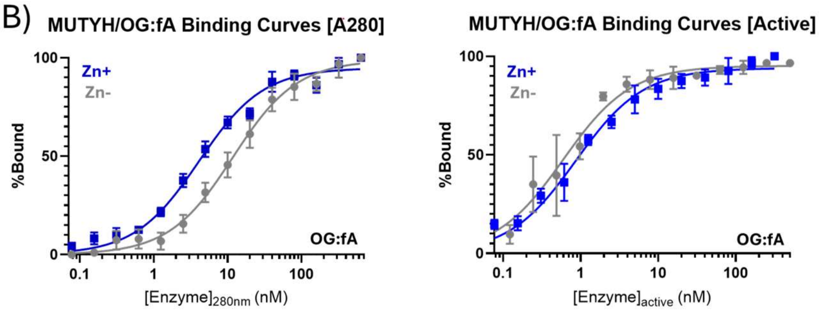


C)

| Enzyme | OG:fA (Absolute) | OG:fA (Active) |
|---|---|---|
| MUTYH (Zn-) | 12 ± 2 | 0.6 ± 0.1 |
| MUTYH (Zn+) | 4 ± 0.4 | 0.8 ± 0.09 |
| $K_{d(Zn-)}/K_{d(Zn+)}$ | 3 ± 0.6 | 0.8 ± 0.2 |

**Figure S7. Zinc-dependent DNA binding affinities of MUTYH measured by electrophoretic mobility shift assay (EMSA). (A)** Representative EMSA gels used to determine equilibrium binding affinities ($K_d$) for MUTYH in the absence (MUTYH(Zn-), top) or presence (MUTYH(Zn+), bottom) of zinc. Binding experiments were performed using either total enzyme concentrations determined by UV absorbance at 280 nm (left) or active enzyme concentrations corrected for the measured active fraction (right). Enzyme concentrations ranged from 0–600 nM (total) or 0–300 nM (active). All assays were conducted with 5 pM 30-bp duplex DNA containing a non-cleavable OG:fA lesion, with the fA-containing strand radiolabeled at the 5′ end using γ-$^{32}$P-ATP. **(B)** Corresponding binding isotherms for MUTYH Zn(+) and Zn(–) at 25 °C, plotted using absolute enzyme concentrations determined by UV–vis absorbance at 280 nm (left) or active enzyme concentrations corrected for enzyme active fraction (right). **(C)** Table of $K_D$ values determined for the Zn- and Zn+ states of MUTYH, and their corresponding ratio under conditions of absolute (UV 280nm) and active concentrations.

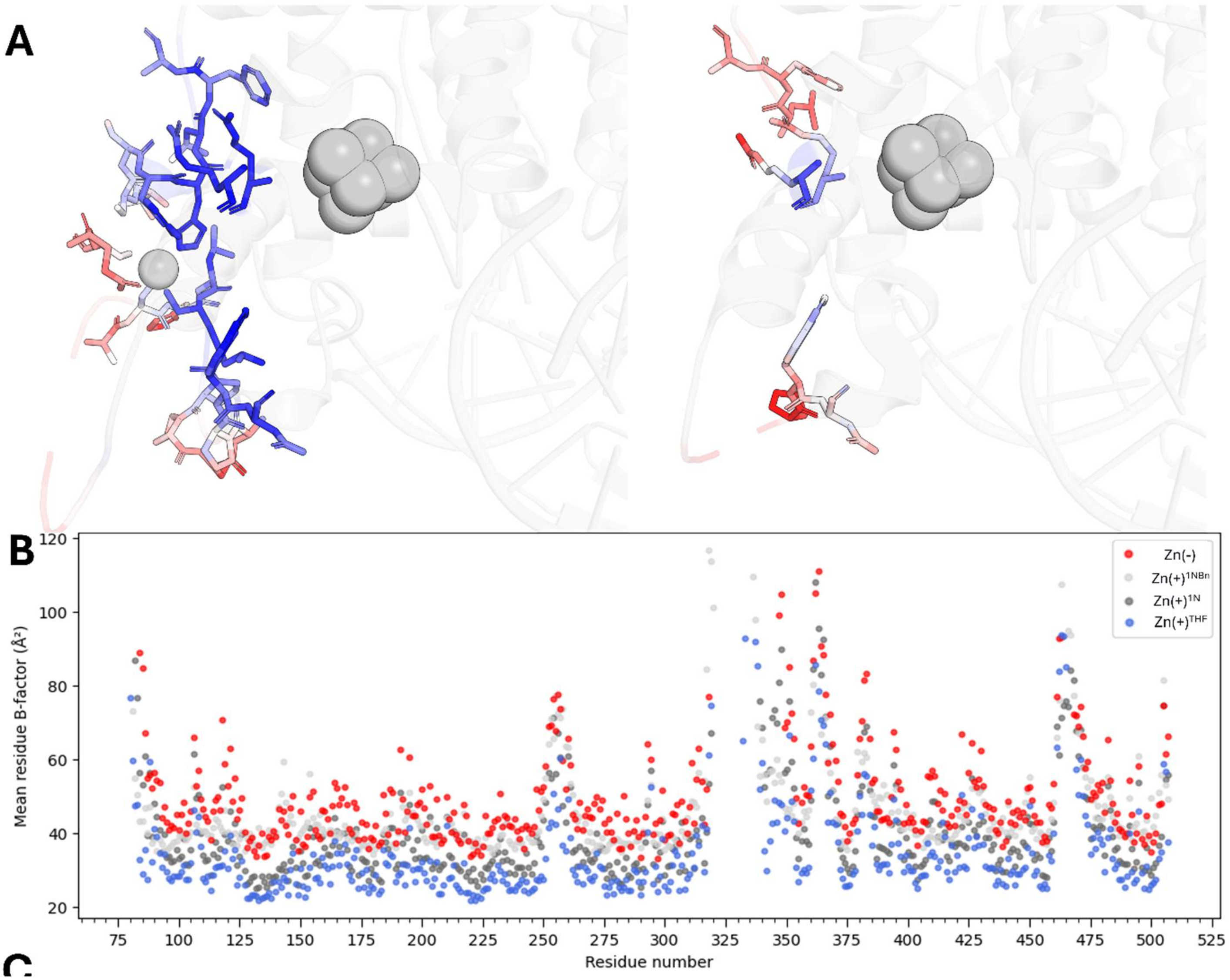


**Figure S8. Residue mobility across MUTYH structures. (A)** Reduced B-factors in residues surrounding the zinc coordination site within the IDC of MUTYH(Zn+), indicative of decreased thermal motion (blue = low B-factor; red = high B-factor), whereas the MUTYH(Zn-) (PDB: 8FAY, (**B**) displays elevated B-factors and increased disorder across the same region. **(C)** Zinc occupancy is associated with reduced side-chain thermal motion in regions proximal to the zinc linchpin motif, with pronounced dampening in the $3_{10}$-helix (aa 80-90) and along the interdomain connector (IDC; aa 321–350). A similar stabilization trend is observed around helix H13 (aa 250–270), including the bridging Arg247 immediately N-terminal to H13. For each structure, per-residue B-factors from the two molecules in the ASU (chains A and D) were first averaged to generate a single composite trace, and this A+D-averaged profile was then self-normalized (z-scored) within each structure prior to overlay plotting.

**Table S2** Composite-region mean B-factors in MUTYH(Zn+) and MUTYH(Zn-)

| ***Composite Region (Residues)*** | ***State*** | ***Mean B-factor (Å²)*** | ***SD (Å²)*** | ***n (Cα atoms)*** |
|---|---|---|---|---|
| ***81–88, 247, 307, 321–350*** | MUTYH(Zn-) | 65.03 | 14.02 | 13 |
| | MUTYH(Zn+) | 49.58 | 14.26 | 33 |
| | **ΔB** | **-15.45** | | |

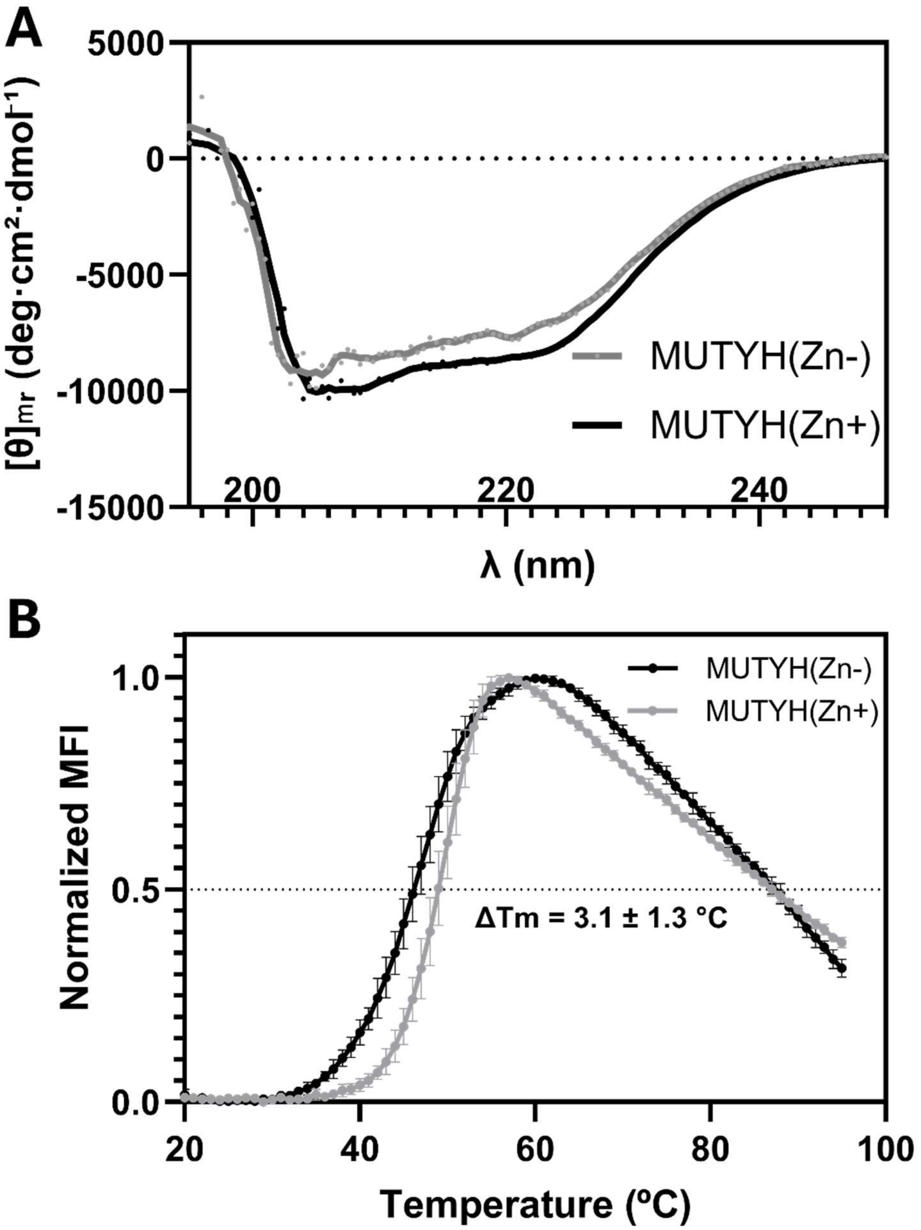


**Figure S9. Effects of $Zn^{2+}$ Binding on MUTYH Thermal Stability and Secondary Structure. (A)** Far-UV circular dichroism (CD) spectra of MUTYH(Zn-) and MUTYH(Zn+) collected on an OLIS CD-20 spectropolarimeter (190–250 nm) at 2 μM protein concentration. Spectra are reported as mean residue ellipticity, $[\theta]$(deg·cm$^2$·dmol$^{-1}$), calculated from the observed ellipticity (mdeg) according to:

$$[\theta] = \frac{\theta_{obs} \times 10^6}{c\, l\, N}$$

where $\theta_{obs}$is the measured ellipticity (mdeg), $c$ is the molar protein concentration, $l$ is the optical pathlength, and $N$ is the number of residues. The MUTYH(Zn+) spectrum exhibits a consistently more negative ellipticity across the far-UV region relative to MUTYH(Zn-), indicative of modestly increased secondary-structure ordering and reduced conformational heterogeneity upon Zn binding. Spectral features below ~205 nm were not recognized due to increased absorbance and reduced signal-to-noise in this region. (***B***) Differential scanning fluorimetry (thermal shift assay) comparing MUTYH(Zn-) and MUTYH(Zn+) reveals that retention of $Zn^{2+}$ stabilizes the enzyme, shifting the melting transition of MUTYH(Zn+) by ~3 °C to higher temperature relative to MUTYH(Zn-).

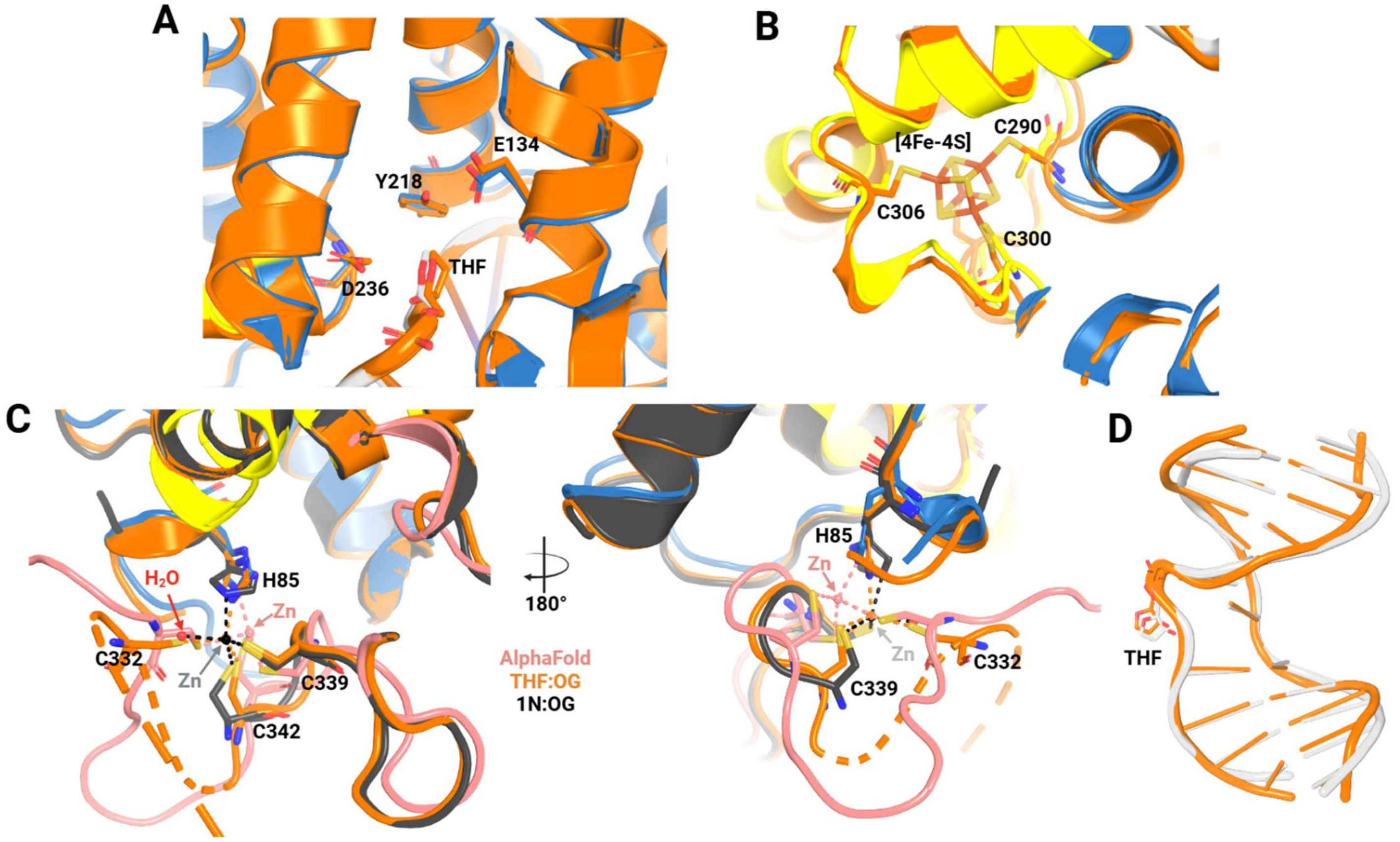


**Figure S10. Structural comparison of the functional component between AlphaFold model and crystallography MUTYH structures.** Structural alignment including the MUTYH portion from the AlphaFold3 model, Product Analog Complex (PAC; THF:OG-containing structure. PDB ID 11IP), orange and, late Transition State Analog Complex (late TSAC; 1N:OG-containing structure. PDB ID 11JI), black. A) Show the comparison of the active site, B) the [4Fe-4S] cluster motif, C) the Zinc Linchpin motif and D) the DNA. For the AlphaFold model the N-terminal domain, the [4Fe-4S] cluster motif IDCL and DNA are shown in blue, yellow, light red and light gray, respectively.
